# Minimisation of surface energy drives apical epithelial organisation and gives rise to Lewis’ law

**DOI:** 10.1101/590729

**Authors:** Marco Kokic, Antonella Iannini, Gema Villa-Fombuena, Fernando Casares, Dagmar Iber

**Affiliations:** Department of Biosystems, Science and Engineering (D-BSSE), ETH Zurich, Mattenstraße 26, 4058 Basel, Switzerland; Swiss Institute of Bioinformatics (SIB), Mattenstraße 26, 4058 Basel, Switzerland; CABD, Universidad Pablo de Olavide, Seville, Spain

**Keywords:** Epithelial Organisation, Lewis’ Law, Surface Energy, Computational Model

## Abstract

The packing of cells in epithelia exhibits striking regularities, regardless of the organism and organ. One of these regularities is expressed in Lewis’ law, which states that the average apical cell area is linearly related to the number of neighbours, such that cells with larger apical area have on average more neighbours. The driving forces behind the almost 100-year old Lewis’ law have remained elusive. We now provide evidence that the observed apical epithelial packing minimizes surface energy at the intercellular apical adhesion belt. Lewis’ law emerges because the apical cell surfaces then assume the most regular polygonal shapes within a contiguous lattice, thus minimising the average perimeter per cell, and thereby surface energy. We predict that the linear Lewis’ law generalizes to a quadratic law if the variability in apical areas is increased beyond what is normally found in epithelia. We confirm this prediction experimentally by generating heterogeneity in cell growth in *Drosophila* epithelia. Our discovery provides a link between epithelial organisation, cell division and growth and has implications for the general understanding of epithelial dynamics.

## INTRODUCTION

The packing of cells in epithelia exhibits striking regularities, regardless of the organism and organ. This suggests that there are fundamental principles to epithelial organization. Epithelial organization has previously been accounted for by topological constraints [1, 2], energy minimization [3–5], and entropy maximization [6]. While each of those theories explains certain aspects of epithelial organisation, a general theory that explains the wide range of epithelial regularities and organizational principles is missing. We will show that epithelial organization can be explained by a minimization of surface energy. Based on our theoretical framework, we explain how cell growth and division as well as mechanical constraints act together to drive epithelial tissue organization.

In epithelia, neighbouring cells adhere tightly to their neighbours via adherence junctions (Fig 1a). As a result, the apical surface of epithelia corresponds to a contiguous, polygonal lattice (Fig. 1a,b, ED Fig. 1a). In epithelia, three edges typically meet in each vertex (Fig 1c). For such a connected planar graph, the total number of cell faces (*f*), cell edges (*e*), and cell vertices (*v*) in the lattice must be related according to Euler’s Formula as *v−e*+*f*=2, and in the limit of large tissues, cells must on average have six neighbours 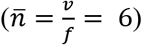 [1, 6]. Even though the frequencies of the different polygon types differ widely between different epithelial tissues, all epithelia analysed to date indeed meet this topological constraint in that their cells have on average close to six neighbours (Fig. 1d,e). This particular property of epithelial organisation thus directly follows from topological constraints.

**Figure 1:**
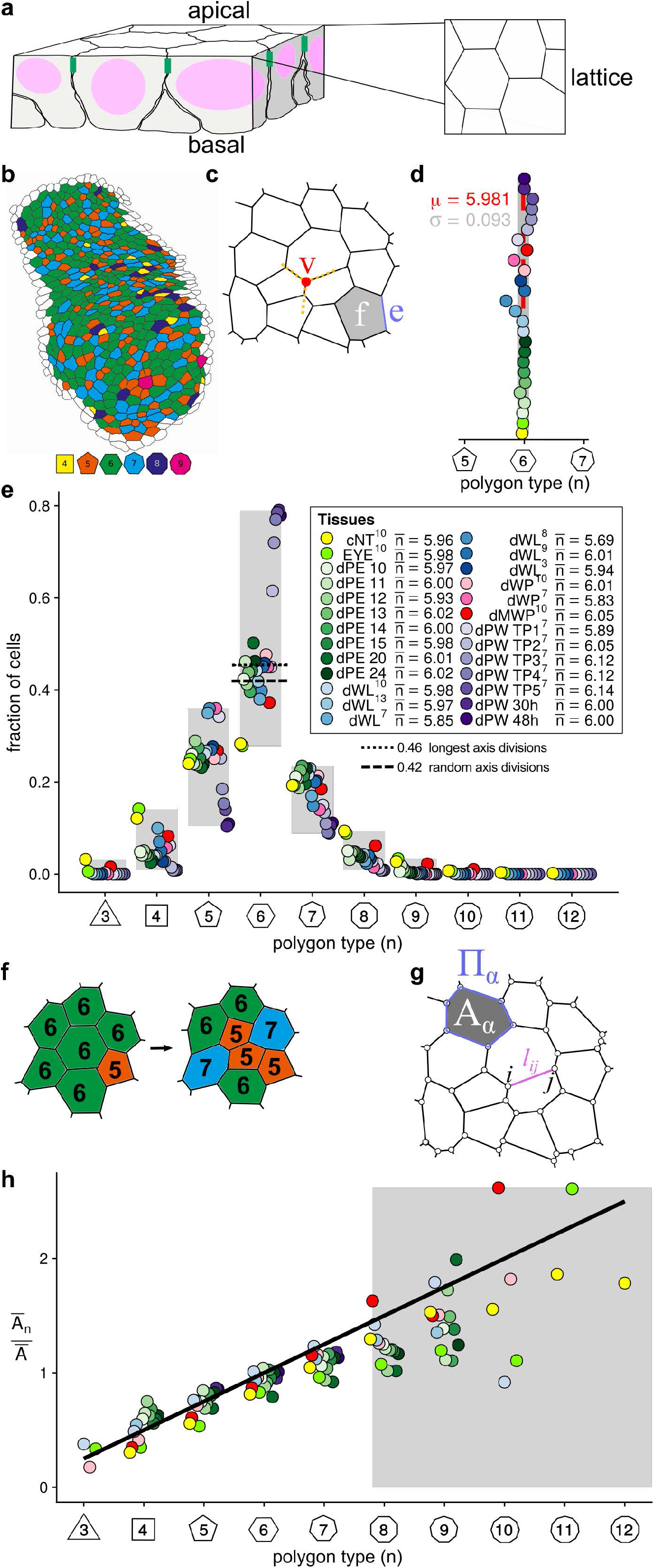
Introduction to Epithelial Packing. **a**, A cartoon of epithelial organisation. For details see text. **b**, A typical epithelial tissue from the *Drosophila* larval eye disc peripodial epithelium (dPE), coloured according to the number of neighbours per cell. The original image with a segmentation mask is shown in ED Fig. 1a. **c**, The apical surface can be represented as a finite, connected, planar graph with *f* faces, *v* vertices, and *e* edges without edge intersection. Three edges are incident to each vertex, i.e. the degree of each vertex is three. **d**, The measured average number of cell neighbours is close to the topological requirement [1, 6], 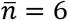, in all tissues; see panel e for the colour code. **e**, Tissues differ widely in the relative frequency of neighbour numbers. Topological models only explain a single (dashed and dotted lines) of these many distributions (shaded range). The dashed line marks the hexagon fraction for randomly positioned cell division axes, the dotted line marks the hexagon fraction when cells are divided perpendicular to their longest axis. The legend provides the measured average number of cell neighbours for each tissue and the references to the primary data [1, 3, 11, 15, 43–45]. Data points for *n*<3 were removed as they must present segmentation artefacts. See ED Fig. 1 for standard deviations and full tissue names. **f**, Topological changes upon cell division have been suggested to result in the measured relative frequency of neighbour numbers. However, topological models only explain a single (panel e, dashed and dotted lines) of these many distributions (panel e, shaded range). **g**, Vertex models are based on finite, connected, planar graphs with vertices *i,j* cell area A_α_ (grey), edge length l_ij_ (blue) and cell perimeter Π_α_ (blue). **h**, The relative average cell area, 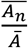, is linearly related to the number of neighbours, *n*, and follows Lewis’ law [6] (Eq. 1, black line) in all reported tissues [15, 44]. The colour code is as in (c), but data is available only for a subset of those tissues. Few cells with more than 8 neighbours have been measured and estimates of the mean areas are therefore unreliable (shaded part).

Topological arguments have also been proposed to explain the particular frequencies of polygon types in the different epithelial tissues (Fig. 1e, ED Fig. 1c). According to those topological arguments the observed frequencies of polygon types are the result of sequential cell division (Fig. 1f) [1, 2]. A key prediction from this topological model is that all epithelial tissues would have the same distribution of polygon type frequencies [1, 2]. The exact frequency would depend on the mode of cell division. However, both in case of a random positioning of the cell division axis (marked by a dashed line in Fig. 1e) as well as in case of cell division perpendicular to the longest axis (Hertwig’s rule [7]) (marked by a dotted line in Fig. 1e), about 45% of all cells would have six neighbours. Contrary to this prediction from the topological model, the frequencies of polygon types vary strongly between tissues, ranging from 30% to 80% hexagons in the different epithelia analysed (Fig. 1e, ED Fig. 1b). Other processes, in addition to cell division, must therefore determine epithelial organisation.

Vertex models explain cell organisation in animal and plant epithelia based on physical arguments [2–5, 8]. Here, the apical surface of each cell is approximated by a polygon. Neighbouring cells share edges such that the epithelium is represented as a polygonal tiling (Fig. 1g). Cells can grow and divide. The mechanical behaviour of the cells is represented either via forces or based on an energy functional. In each time step, the vertices of cells are moved based on the equations of motion until the forces balance at each vertex, which corresponds to the energy minimum. As the forces acting at each vertex can be derived from the energy functional, the two approaches are equivalent, even though implementations may differ [9].

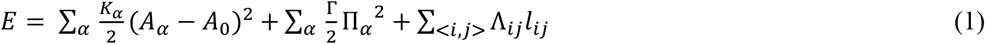

represents a well-studied energy functional for epithelial dynamics [3, 5]. Here, the first term serves to limit the area variability in the tissue, the second term describes cell contractility, and the last term represents the net effect of cell adhesion and surface (line) tension [3, 5]. In the first term, *K_α_* ≥ 0 is a constant, the so-called area contractility, *A*_0_ is the target apical area, and *A_α_* is the apical area of cell α; the sum goes over all cells in the tissue. In the second term, Γ is the contractility, and Π_*α*_ is the perimeter of cell α; the sum goes over all cells in the tissue. Finally, in the last term, Λ_*ij*_ is the surface (line) tension and *l_ij_* is the length of the edge between vertex *i* and *j*; the sum goes over all edges in the tissue. The wide range of measured polygon type frequencies can be reproduced by adjusting the mechanical parameters Λ_*ij*_ and Γ relative to *K_α_* [3]. In addition to the mechanical parameters, also the mode of cell division affects the polygon type frequencies [2]. It has therefore been argued that both topological and mechanical effects determine epithelial organisation in the vertex model [2].

In 1928, F.T. Lewis documented a further intriguing property of epithelia [10]: the average apical area, 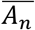, of cells with *n* neighbours is linearly related to the number of neighbours *n*, such that

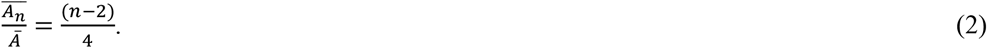

Here, *Ā* refers to the average apical cell area in the tissue. All analysed epithelial tissues follow Lewis’ law (Fig. 1h, ED Fig. 1c) despite the different frequencies of neighbour numbers in the different tissues (Fig. 1e, ED Fig. 1b). Vertex models can reproduce Lewis’ law, but only for a small subset of the parameter space (spanned by the relative contractility, 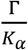, and relative line tension, 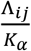) [3, 5, 8], and it is unclear what restricts epithelia to all fall within this narrow parameter range. Similarly, Voronoi tessellations can give rise both to the measured range in the neighbour frequencies and to Lewis’ law, but again not for all Voronoi tessellation paths, and Voronoi tessellation paths do not necessarily recapitulate cell division planes [11]. Finally, entropy maximisation rather than energy minimisation has been proposed as a driving force for Lewis’ law [6], but this has since been ruled out [12, 13]. In conclusion, none of the current theories can explain the robust emergence of Lewis’ law in animal and plant epithelia analysed to date, suggesting that an important driving force of epithelial organisation has so far been overlooked. So, what is the underlying principle that drives all epithelia to organise according to Lewis’ law?

In the following, we show that the observed epithelial order indeed minimises energy. The key aspect that had been overlooked so far is the impact of the variability in the apical cell area on epithelial organisation. As a result of cell growth and division, apical areas differ in epithelia. In that case, a tiling with a mix of polygon types (rather than hexagonal tiling) has the smallest combined perimeter. Lewis’ law emerges because the side lengths in the apical polygonal tiling are then the most similar in length, resulting in the most regular polygonal shapes within a contiguous lattice of cells with different apical areas. Regular polygons result in the smallest cell perimeter per area, and regular polygonal lattices therefore minimize the contact surface area between cells, and thus the contact surface energy. Interestingly, our theory predicts that the relationship between cell area and neighbor number follows a quadratic law, rather than the linear law found by Lewis. Simulations predict that this quadratic relationship can only be observed in the limit of much higher apical area variability than what is normally found in epithelia. We tested the theory by generating such higher apical area variability in the *Drosophila* wing disc epithelium, and we indeed confirm the predicted quadratic scaling law. Moreover, we predict and confirm experimentally that the fraction of hexagons in a tissue depends on the area variability in the tissue. We conclude that the observed epithelial organisation is driven by a minimisation of the lateral contact surface energy between epithelial cells. Epithelial organisation and Lewis’ law can thus be understood by considering only two variables: tension and the variability of the apical cell areas in the epithelial tissue. Growth and cell division affect the epithelial organisation via their impact on the apical cell areas.

## RESULTS

### Epithelial Organisation when all apical areas are equal

We seek to understand the observed distribution of neighbour numbers and the emergence of Lewis’ law in epithelia. Let us first consider a tissue with equal line tension in all edges, Λ_*ij*_ = Λ, and equal apical areas, i.e. *A_α_* = *A*_0_. The energy *E* (Eq. 1) then only depends on the combined cell perimeter in the tissue, Π = Σ_*α*_ Π_*α*_ = 2Σ_<*i,j*>_ *l_ij_* the factor 2 comes in as neighbouring cells share edges in the vertex model. For a given area, the perimeter of a polygon depends on its shape and on its number of neighbours: regular polygons have the smallest perimeter per area, and the polygon perimeter *Π* declines non-linearly as the number of vertices, *n*, increases (Fig. 2a),

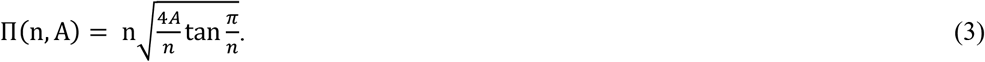

**Figure 2:**
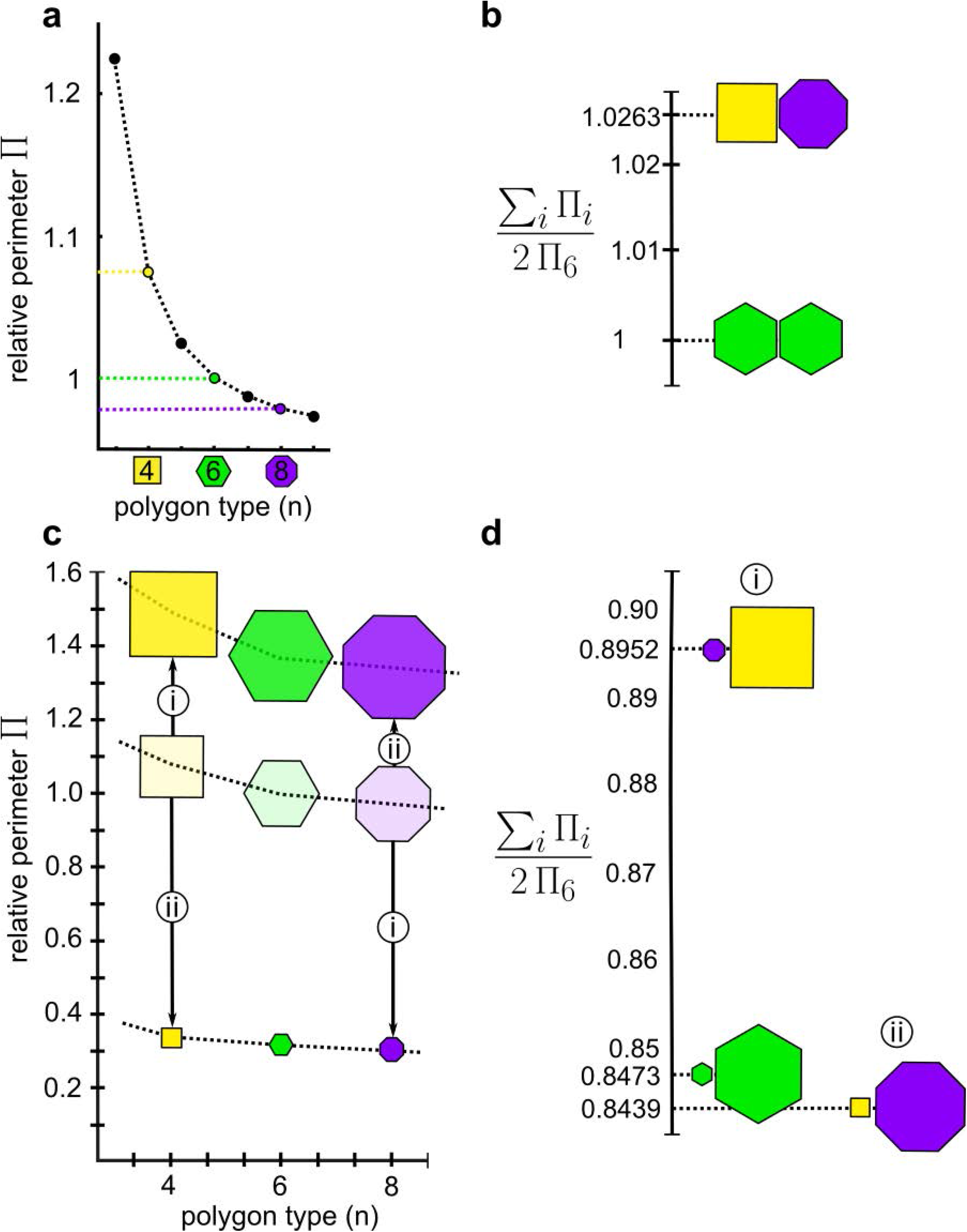
The impact of apical area variability on Epithelial Order. **a**, The perimeter Π of regular polygons declines non-linearly with increasing vertex number, *n*. **b**, In contiguous lattices composed of polygons with three edges per vertex, the average number of vertices must be six. The combined perimeter of non-hexagonal polygons with on average six vertices, i.e. a square and an octagon, is larger than that of two hexagons. **c**, The perimeter of regular polygons dependent on the vertex number, *n*, for different areas, A = 0.1, 1, 1.9. **d**, The impact of area variability. Two hexagons that differ in their area (*A* = 0.1 and *A* = 1.9) have a lower combined perimeter than two hexagons that have the same area (*A* = 1). Similarly, the combined perimeter of the square and the octagon is smaller when the two polygons differ in their area (A = 0.1 and A = 1.9 instead of A = 1 for both polygons). However, the square and the octagon have a smaller combined perimeter than the two unequally sized hexagons only if the smaller polygon (A=0.1) has fewer vertices (square) and the larger polygon (A=1.9) has more vertices (octagon) (case ii). The combined areas are identical in all cases; the combined perimeter is normalised with the combined perimeter of two hexagons that have the same area.

The perimeter per area is smaller, the larger the number of vertices. However, in contiguous tissues with three edges per vertex, cells must respect the topological constraint 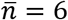 (Fig. 1c, d), i.e. cells must have on average six neighbours [1, 6]. Therefore, if some cells have more than six neighbours, other cells must have less than six neighbours. As can be seen in Figure 2a, for a given area, cells with fewer than six vertices gain more in perimeter than what the cells with more than six vertices lose in perimeter. For instance, a square and an octagon, which have on average six vertices, have a larger combined perimeter than two hexagons (Fig. 2a,b). Given the nonlinear dependency of the perimeter on the number of vertices (Fig. 2a), hexagonal tiling always has a smaller combined perimeter than any mixed polygon tiling, and purely hexagonal tiling is then the energetically most favourable configuration. Why would tissues deviate from hexagonal packing and follow Lewis’ law?

### The impact of Apical Area Variability on Epithelial Order

The key aspect is the variability in the apical cell areas. As a consequence of growth, cell division, and nuclear movement during the cell cycle, apical cell areas in epithelia differ widely between cells. Changes in the polygon area shift the perimeter curve in a nonlinear fashion (Fig. 2c). As a result, if we again compare the two hexagons with a square and an octagon, we notice that differences in cell area reduce the combined perimeter for all configurations, even though the combined areas of the polygons is the same as before (Fig. 2d). The strongest reduction in the combined perimeter is observed when the smaller polygon has fewer vertices (square) and the larger polygon has more vertices (octagon) (Fig. 2c,d, case ii). Given the relation between polygon perimeter, area, and neighbour number (Eq. 3), this observation holds not only for this particular example, but for any combination of polygons that has on average six vertices (Euler’s formula). While the smaller combined perimeter explains why smaller cells have on average less than six neighbours and larger cells have on average more than six neighbours, it does not yet explain why the linear Lewis’ law emerges.

### Lewis’ Law minimizes Surface Energy by driving polygons to be regular

In our mathematical analysis, we have so far assumed that all polygons are regular, which implies that they are equilateral and equiangular. Regular polygons have the smallest possible perimeter at a given apical cell area, and energy minimisation should therefore drive polygonal lattices to the most regular polygonal shapes. However, to fit within a contiguous lattice, neighbouring cells need to have the same side lengths (Fig. 3a). If all cells had the same apical area (Fig. 3b, blue line), then cells with fewer vertices would have larger average side lengths (Fig. 3c, blue line). Cell edges would thus need to be stretched to constitute a contiguous lattice (Fig. 3a). If the apical cell areas follow Lewis’ law, then the differences in side lengths are much smaller between different polygon types (Fig. 3c,d, black lines). In fact, equal side lengths,

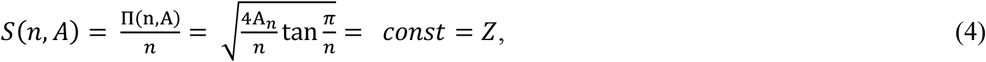

and thus regular polygons would be obtained (Fig. 3c,d, yellow lines) if

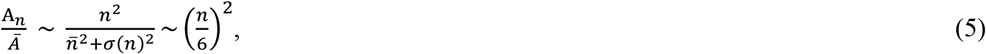

where 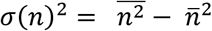 is the variance in the cell neighbour numbers. Here, we solved Eq. 4 for *A_n_*, i.e. 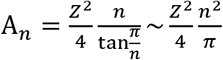, and estimated the average area as 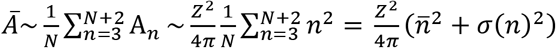, where *N* is the highest observed cell neighbour number. Using Euler’s formula, 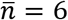, we then obtain Eq. 5, which is very similar to Lewis’ law, except that the relative areas depend quadratically rather than linearly on the neighbour numbers. So, why do epithelia follow the linear Lewis’ law (Eq. 2) rather than the quadratic relation in Eq. 5?

**Figure 3:**
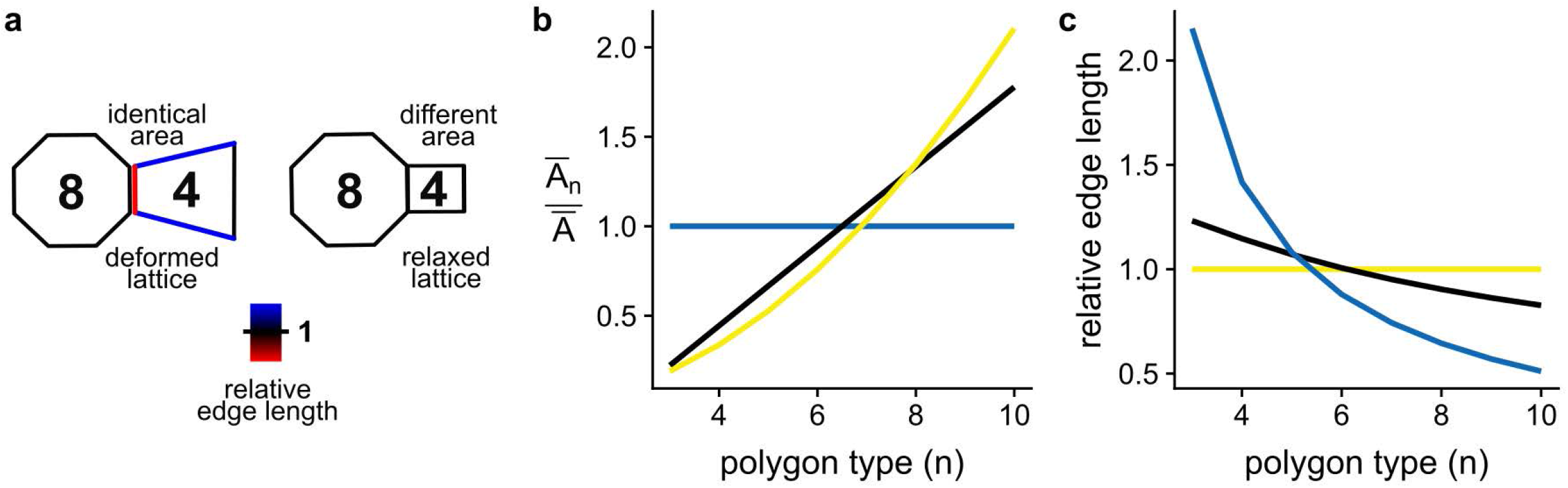
Lewis’ law reduces variability in side length. **a**, To fit into a contiguous lattice, cells must have equal side lengths or be stretched. **b,c** If all cells have the same apical area (b, blue line), then the edge length changes with the polygon type (c, blue line). If the cell areas follow Lewis’ law (Eq. 2), then the change in edge length is much smaller (black lines). All edges have the same length, if the polygon type-dependency of the apical areas follows Eq. 4 (yellow curves).

### LBIBCell Simulations of Epithelial Tissue Dynamics predict a critical role of the apical area variability for Lewis’ law

To address this question, we need to use a computational framework that permits cells to deform within the tissue, and we used the simulation framework `LBIBCell` [14] to simulate the apical cell dynamics in a 2D plane with high geometric resolution (Fig. 1a, Fig. S1, Movie S1). The opposing effects of cell-cell adhesion and lateral surface tension are represented by springs at cell-cell junctions, and the effect of cortical tension is represented by springs along the cell surface (Fig. 4a). The most quantitative data is available for the *Drosophila* wing disc epithelium (or disc), and all parameter values could be inferred from available quantitative data (see Supplementary Material for details). Thus, the fraction of hexagons depends on the two mechanical parameters (Fig. 4b), and we adjusted the boundary tension and cell-cell junction strength to reproduce the frequencies of cell neighbours measured by Heller and co-workers in *Drosophila* wing discs [15] (Fig. 4c). We used the measured declining growth rate [16, 17] to reproduce the measured wildtype wing disc growth kinetics [18] (ED Fig. 2a,b), and we set the cell division threshold (Fig. 4d) such that we reproduce the measured average apical cell area (5.34 μm^2^) [15] (Fig. 4e) and the fraction of mitotic cells [5] (ED Fig. 2c). Cells were divided perpendicular to their longest axis (Hertwig’s rule [7]), as observed in wing discs [2, 19], but randomised cell division angles give similar results (ED Fig. 3). A fixed cell division threshold (Fig. 4d) results in a much narrower area distribution (CV = std/mean = 0.21) than measured (CV = 0.38, Fig. 4e) and the resulting slope in the Lewis’ plot is shallower than observed (Fig. 4f). Higher area CV values can result from less precise cell division thresholds, asymmetric cell division, and/or cell deformation, i.e. changes in apical area without change in cell volume as occurs in apical constriction. We randomised the critical size for cell division (Fig. 4g, ED Fig. 4, Supplementary Material) to reproduce the measured apical area distribution (Fig. 4h). For the measured apical area CV, Lewis’ law emerges in the simulations (Fig. 4i), thereby confirming the key importance of the apical area variability for epithelial packing. When we further increased the apical area variability (Fig. 4j,k), the simulations diverted from Lewis’ law (Fig. 4l, black line) and matched the predicted quadratic curve (Eq. 4) (Fig. 4l, yellow line). The simulations show and thus predict that the optimal average area for each polygon type is obtained only with wider apical cell area variability than normally observed in epithelia.

**Figure 4:**
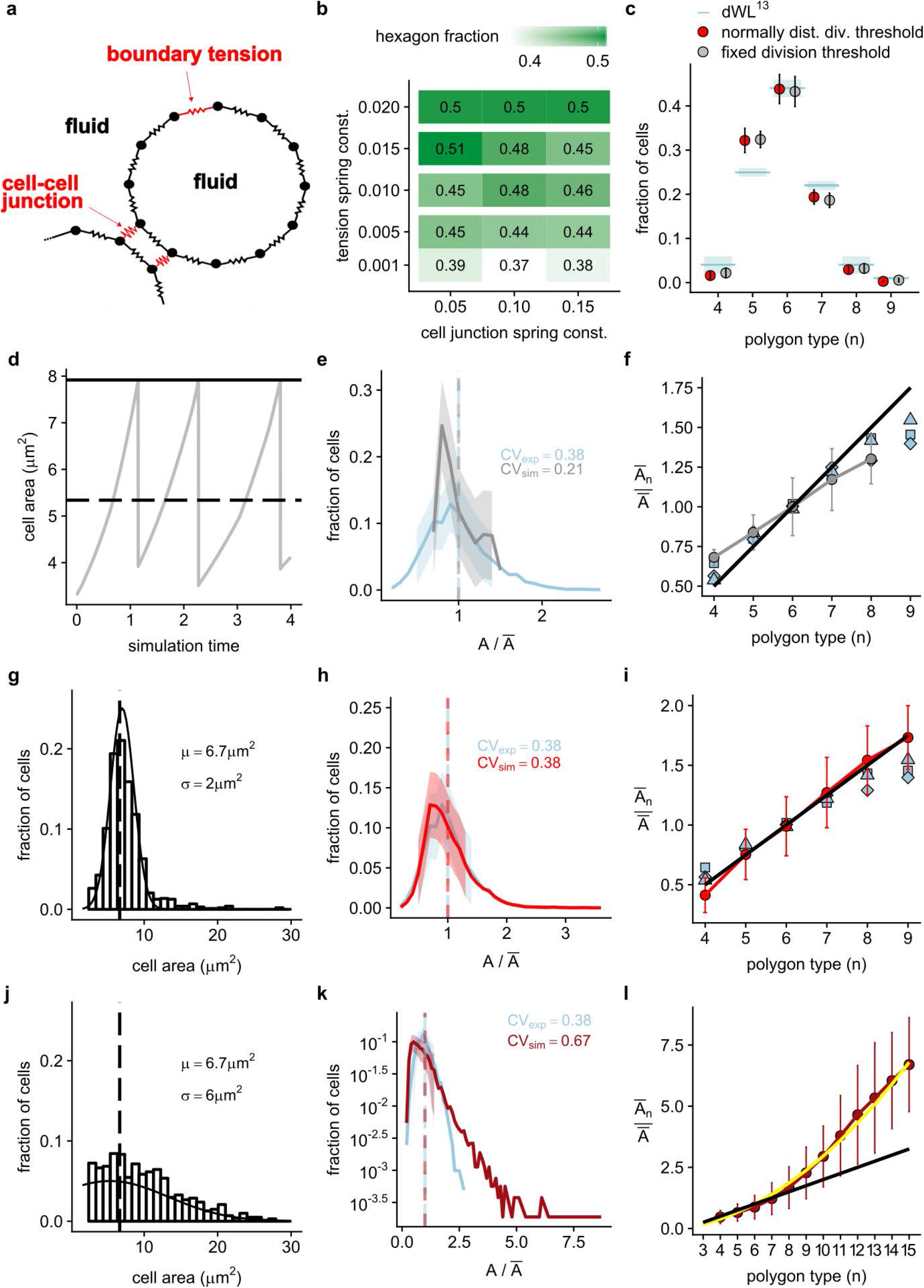
Simulations in LBIBCell confirm the impact of the cell size distribution on epithelial packing. **a**, Representation of tissue and cells in `LBIBCell` [14]. Cortical tension and cell-cell adhesion are incorporated via springs (red). **b**, The fraction of hexagons increases with higher membrane tension and stronger cell adhesion in the simulations. **c**, The measured hexagon frequency in the wing disc [15] (blue line) can be reproduced. **d-f**, Cell division at a fixed cell size (d) results in a narrow cell size distribution (e), and fails to recapitulate Lewis’ law (Eq. 2, black line) (f). **g-i**, With a normally distributed cell division threshold (with mean μ and standard deviation c) (g) the simulations (red) recapitulate the measured (blue) cell size distribution [15] (h) and Lewis’ law (black line) (i). The blue symbols in f,i represent three independent measurements [15]. **j-l**, With an even wider distribution of cell division thresholds (j) the cell size distribution widens further (k) and the simulations deviate from Lewis’ law (black line) and approach the curve for which all side lengths are equal (yellow line, Fig. 3b) (l).

### Experimental Test: higher apical area variability results in the predicted quadratic law

We sought to test this prediction by artificially creating epithelia with wider apical cell area distributions. To this end, we induced the expression of an RNAi targeting the growth regulator *gigas/TSC2* [20] in cell patches randomly located within *Drosophila* wing discs. *gigas* attenuation results in larger cells because cells repeat S phase without entering M phase [20]. Indeed, in mosaic *gigas/TCS*-RNAi wing discs we observe cells with substantially larger apical areas than in the control, scattered throughout the epithelium (Fig. 5a). Importantly, most *gigas/TSC2* cells still have an apical area in the wt range, but the largest apical areas are substantially larger than in the control such that the area distribution is considerably wider (Fig. 5b).

**Figure 5:**
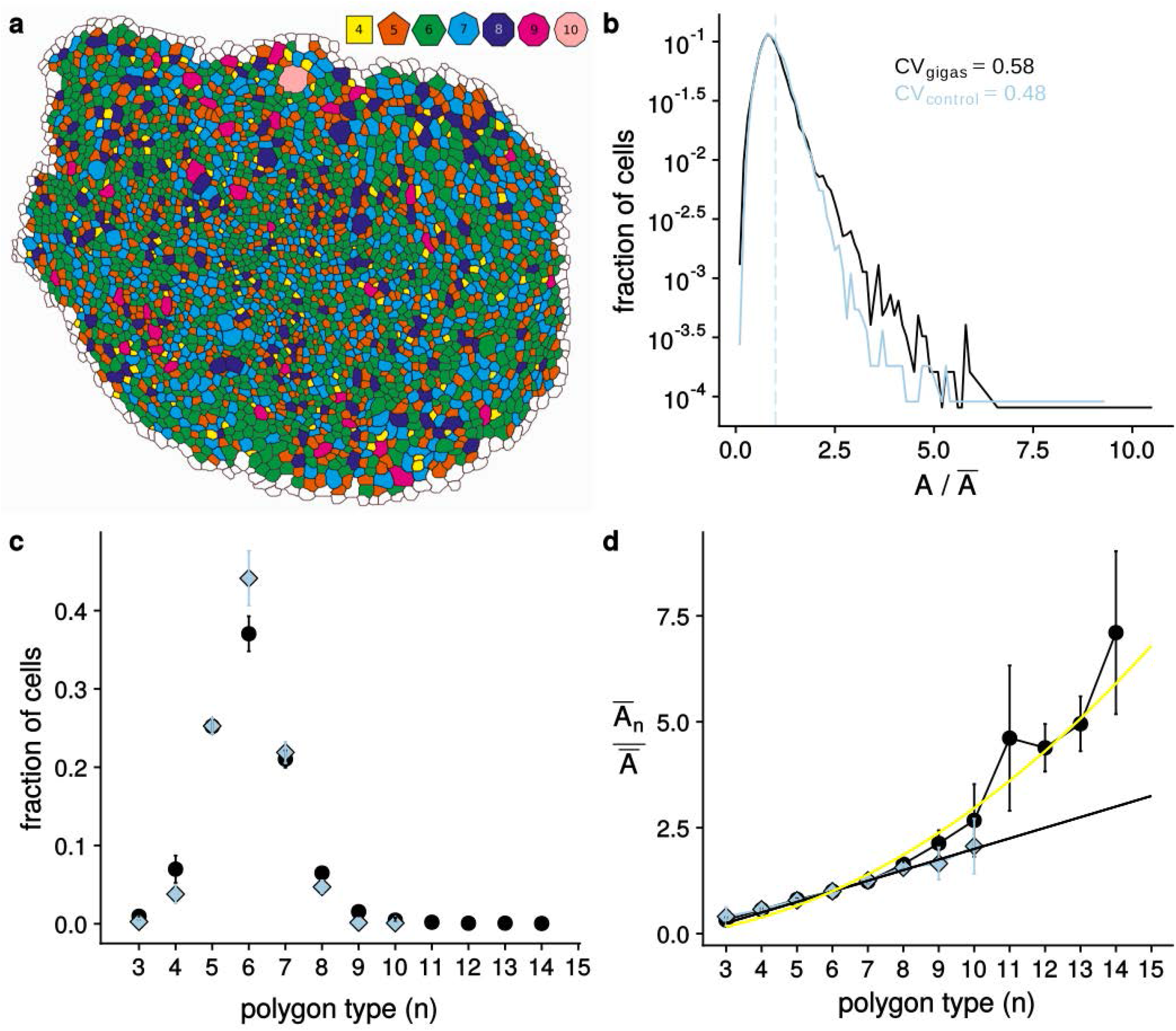
Experimental Confirmation of the generalised Lewis’ law. **a**, Representative epithelial tissue from a wing disc with *gigas* RNAi clones. **b**, The apical cell area distribution, **c**, the relative frequency of neighbour numbers, and **d**, the relative average apical area, 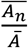 for different neighbour numbers in control (blue) and mutant (“gigas”, black) wing discs. The black line in panel d represents Lewis’ law (Eq. 2); the yellow line the quadratic relation (Eq. 5).

Based on our theory, we make two predictions: first, that the largest cells will on average have more neighbours than any of the wt cells – and indeed in the gigas/TCS-RNAi wing discs the polygon distribution is wider and cells with more than 10 neighbours are observed (Fig. 5c). Consistent with theoretical considerations (Fig. 2c,d) and our simulations (ED Fig. 4c), not only the polygon distribution is wider, but also the hexagon frequency is lower (Fig. 5c). The second prediction from the model and the simulations is that the tissue with the wider area distribution will follow a quadratic relationship rather than the linear Lewis’ law (Fig. 5d). Indeed, the average apical cell area for each neighbour number now corresponds to the predicted optimum (Eq. 4) – that is, a quadratic relationship, rather than the Lewis’ law (Fig. 5d).

### The apical area distribution defines the hexagon fraction

Finally, based on our theory, we can predict and experimentally test a novel relationship. Lewis’ law (Eq. 2; Fig. 6a, black) and the quadratic law (Eq. 5; Fig. 6a, yellow) relate the apical area to a polygon type (Fig. 6a, dotted lines). Based on these laws, we expect that the wider the area distribution, the smaller the fraction of cells that falls within the area range that is predicted to form hexagons (Fig. 6b, shaded green). This trend is indeed observed in our experiments in the *Drosophila* wing disc (Fig. 5b,c). We can make this relationship more precise and quantitatively test it with the available data.

**Figure 6:**
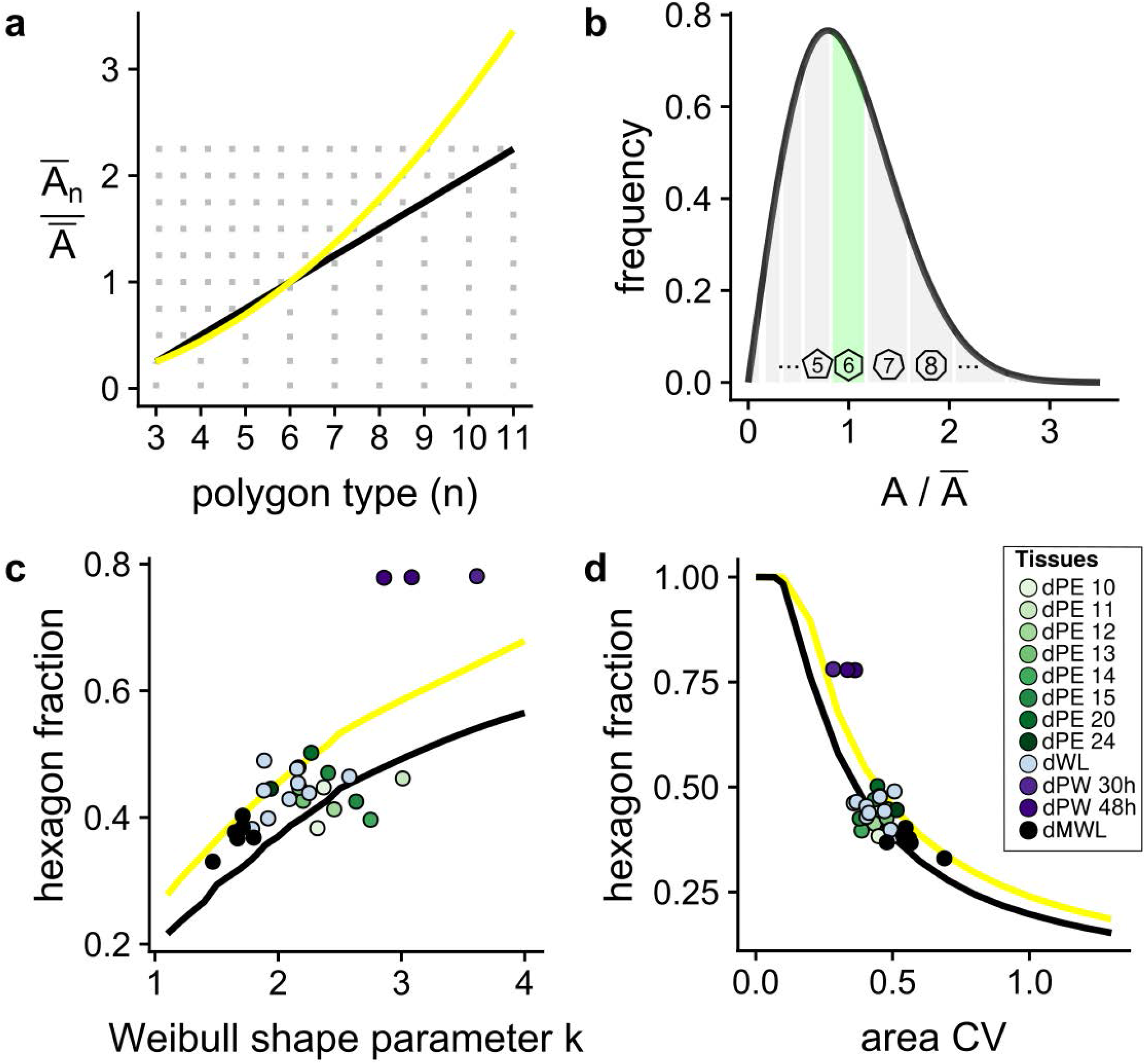
The apical area distribution defines the hexagon fraction. **a**, Lewis’ law (Eq. 2, black line) and the quadratic law (Eq. 5, yellow line) relate the relative apical area to a polygon type *n* (dotted lines). **b**, Weibull distribution (Eq. 6) with *λ* = 1.12, *k* = 2 to approximate the normalised area distribution. The white lines indicate the limits of the polygon types, such that the average area for each polygon type *n* is closest to 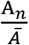. The area shaded in green indicates the part of the area distribution that would correspond to hexagons. **c,d**, The fraction of hexagons for (c) different Weibull shape parameters *k*, and (d) different area CV values (Gaussian distribution) as predicted by the linear Lewis’ law (Eq. 2, black line) or the quadratic law (Eq. 3, yellow line) and for the data (dots). The abbreviations are as in ED Figure 1. dMWL refers to the wing disc with *gigas* RNAi clones from Figure 5.

To predict how the fraction of hexagons depends on the width of the area distribution, we approximate the normalised area distributions either by a Gaussian distribution with variance *σ*^2^,

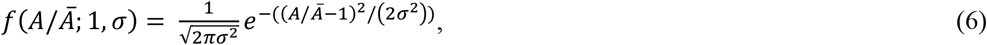

or by a Weibull distribution, which is positive for positive values of *A/Ā*, but is zero otherwise,

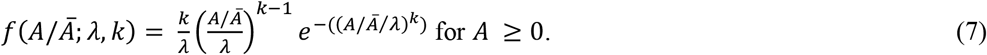

Here, *λ* is the scale parameter that predominantly affects the position of the maximum, and *k* > 1 is the shape parameter of the Weibull distribution that predominantly affects the width of the distribution; as *k* increases, the distribution sharpens (ED Figure 5). The expected fraction of hexagons can then be derived by first assigning each polygon type *n* to a part of the normalised area distribution such that the average area for each polygon type *n* is closest to 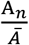 (Fig. 6b, white lines), and by subsequently calculating the value of the cumulative distribution function for hexagons (Fig. 6b, shaded green). The predicted fraction of hexagons for the different Weibull shape parameters *k* and for different area CV values (Gaussian distribution) (Fig. 6c,d, lines) fit the data (Fig. 6c,d, dots) very well, both when we use the linear Lewis’ law (Eq. 2, black line) or the quadratic law (Eq. 3, yellow line). A hexagon fraction below these predicted values arises when cells seek to maximise rather than to minimise their lateral cell-cell contact surface, as is the case when the adhesion strength is increased in the `LBIBCell` simulations (ED Figure 6).

In summary, the theory, computational modelling, and experimentation indicate that the minimisation of surface energy drives the emergence of Lewis’ law, and that a generalization of this law, that applies in the limit of a broader variability in apical cell areas than normally found, is described as a quadratic relationship between average cells area and neighbour number.

## DISCUSSION

Different theories have been proposed to explain epithelial order [1–6, 11]. We have now shown that epithelial organisation can be understood in terms of energy minimisation (Fig. 7). Lewis’ law emerges in all epithelial tissues studied to date because of the particular role played by apical cell area variability, which we describe here. Growth, cell division, and nuclear migration affect epithelial order indirectly via their impact on the variability in the apical cell area.

**Figure 7:**
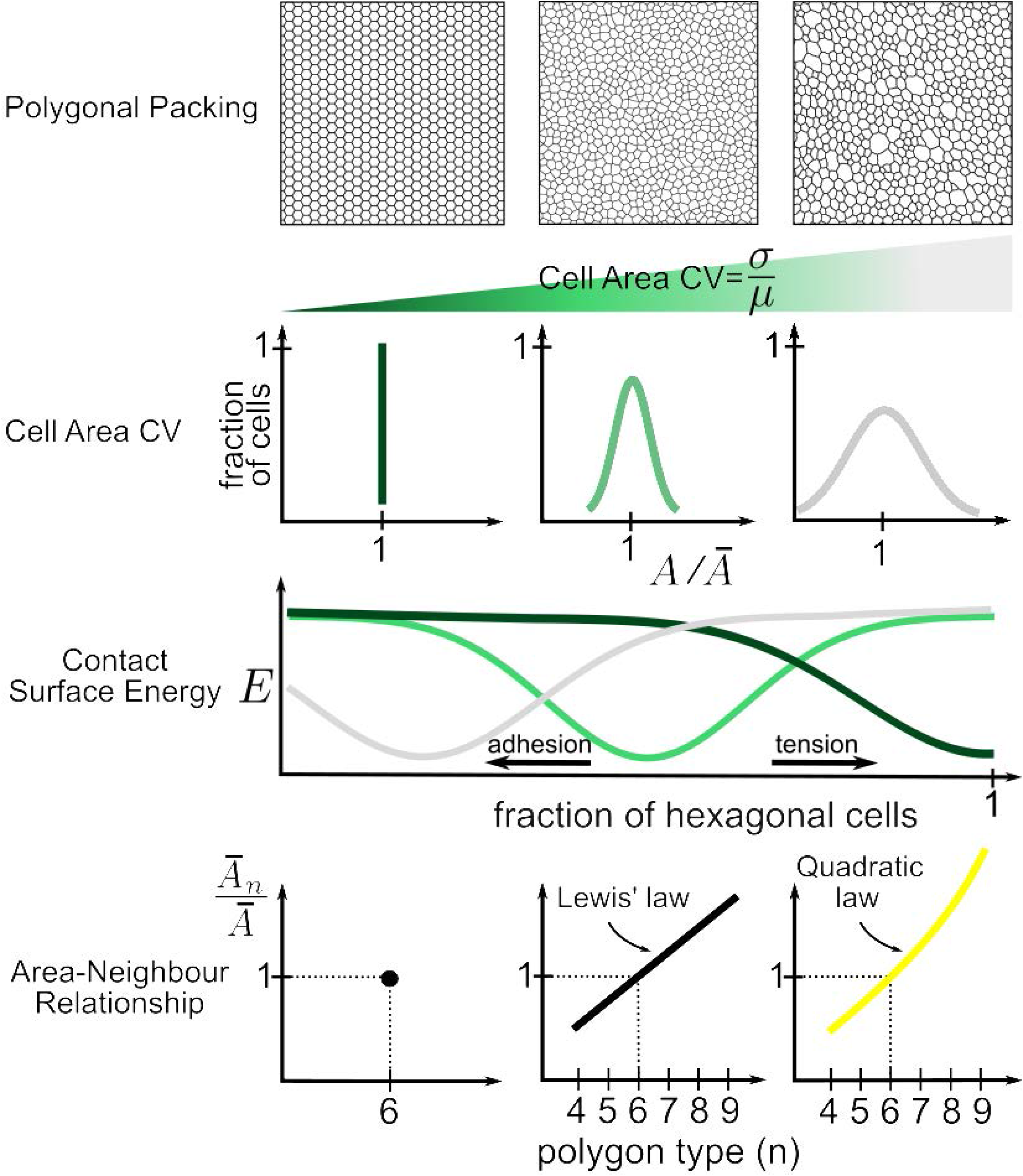
Summary Figure. **(Left column)** If all cells have the same apical area, then hexagonal packing results in the smallest average perimeter per cell, and thus in the lowest possible surface energy. **(middle column)** For an intermediate width of the apical area distribution, lowest surface energy is achieved for non-hexagonal packing and if the average areas per polygon type follow Lewis’ law (Eq. 2). **(right column)** For wide apical area distributions, the lowest surface energy is achieved for non-hexagonal packing and if the average areas per polygon type follow a quadratic dependency on the number of neighbours (polygon type) (Eq. 4). In this case, the side lengths of all polygons are the most similar, and cells thus do not need to be stretched to fit into a contiguous lattice.

We derived Lewis’ law as the configuration that minimises the overall cell perimeter in a contiguous tissue. Cells in a contiguous tissue seek to minimise their perimeter when the effects of cell contractility and lateral surface tension dominate the effects of cell-cell adhesion. In a tissue with equal apical areas, hexagonal tiling results in the smallest combined perimeter (Fig. 2a,b) and is therefore the configuration with the lowest energy (Fig. 7, left column). This configuration has previously been referred to as ground state [3]. In real tissue, apical areas differ as a result of growth, cell division, and interkinetic nuclear movements. If the variability is sufficiently high, cell lattices composed of mixed polygon types have a smaller combined perimeter if the smaller polygon has fewer than six and the larger polygon has more than six neighbours (Fig. 2c,d). Overall, the average number of neighbours must be six due to topological constraints (Euler’s formula) [1, 6]. Lewis’ law (Eq. 2) emerges because cells then have a more similar side length (Fig. 7, middle column), and regular, equilateral polygons have the smallest perimeter at a given area. Equal side lengths of all cells require a quadratic (Eq. 5) rather than a linear (Eq. 2) Lewis’ law. We predict and confirm experimentally that such quadratic law only emerges at higher apical area variability than previously observed in epithelia (Fig. 7, right column).

The role of apical area variability and the regularity of polygons is also observed in other simulation frameworks. Thus, Voronoi tessellations result in epithelia-like packing [11, 21] if the employed algorithm produces nearly equilateral polygons. In vertex models, Lewis’ law has previously been observed only when the mechanical parameters are sufficiently high compared to the area contractility [3]. This can now be understood in terms of our results: a small area contractility corresponds to a weak constraint on the area variability (Eq. 1), which allows a sufficiently wide area distribution for Lewis’ law to emerge. If the mechanical parameter values are further increased compared to the area contractility, then the linear Lewis’ law disappears in the vertex models [3]. Based on our analysis, we expect that a quadratic relationship should be observed in the vertex model when the area contractility is sufficiently small compared to the mechanical parameters.

According to Lewis’ law and the quadratic law, the area distribution determines the width of the neighbour number distribution and thus the fraction of hexagons (Fig. 6). So how do the mechanical properties of cells affect epithelial organisation? The critical aspect here is that in Eq. 1 the line tension term is linear while the contractility term is quadratic in the perimeter. If both constants are positive, then cells will always seek to minimise their combined perimeter, and the area distribution will determine epithelial packing independent of the exact value of the mechanical parameters. Likewise, if both constants are negative, cells will seek to maximise their combined perimeter and will assume deformed cell shapes; these latter tissues are predicted not to follow Lewis’ law. Here, we note that, despite their shape, puzzle cells in plants do not fall into the latter category. This is because puzzle cells do not emerge to maximise their cell perimeter, but to minimize stress in large cells [22]. Importantly, the puzzle shape emerges only after cells have stopped dividing and a contiguous lattice with defined neighbour numbers has already formed, which can no longer change because of rigid cell walls [22]. Accordingly, puzzle cells in plants still follow Lewis’ law [23].

Epithelial organisation reflects the mechanical properties of cells only if the line tension constant, Λ_*ij*_, and contractility, Γ, have different signs. The line tension constant will typically be negative because the effects of cell-cell adhesion can be expected to be stronger than those of lateral surface tension; contractility will be positive due to the effects of the acto-myosin ring. Given that the linear term is then negative and the quadratic term is positive, a minimal cell perimeter will minimise the energy (Eq. 1) only for those cells that have a cell perimeter above a critical value, and this critical value depends on the relative value of the negative line (surface) tension constant, Λ_*ij*_, and the positive contractility, Γ. Accordingly, an increase in the cell-cell adhesion strength relative to cell contractility results in deformed cells (ED Fig. 6a-d), a smaller fraction of hexagons (ED Fig. 6e), and a deviation from Lewis’ law (ED Fig. 6f). Likewise, external forces that stretch the entire tissue result in a deviation from Lewis’ law (ED Fig. 3d). As a practical consequence, relative forces (and thus stresses) in an epithelium can be inferred from the observed cell shapes [24, 25].

We conclude that Lewis’ law emerges as cells seek to minimize the overall cell perimeter in a tissue, which is the case when contractility and surface tension dominate the effects of cell-cell adhesion. The mechanical properties of the cells determine to what extent cells seek to minimise their perimeter – and thus to what extent Lewis’ law applies in a tissue. The fact that Lewis’ law is observed throughout epithelia shows that the combined effects of contractility and surface tension typically dominate the effects of cell-cell adhesion and cells thus seek to minimise their perimeter while forming a contiguous lattice. Whether the linear (Eq. 2) or quadratic (Eq. 5)

Lewis’ law is observed, depends on the area variability.

Epithelial cells interact not only at the apical side, but also along the apical-basal axis. Accordingly, similar physical constraints are likely to apply. Further quantitative studies are required to elucidate the 3D organisation of epithelia.

## METHODS

### Imaging of Tissue Samples

*Drosophila* (wild type Oregon-R strain) eye discs and wing discs were dissected from third instar larvae or pupae. Eye discs were staged according to their number of ommatidial rows (as this number increases as development proceeds). Pupal wing discs were dissected at 30 and 48 hours post puparium formation. Staining and confocal imaging was carried out as described[26]. Primary antibodies used were mouse anti-Dlg (4F3 Hybridoma bank, 1:500; eye discs) and rabbit anti-aPKC (Abcam, AB5813, 1:500; pupal wing discs).

Mosaic induction of *gigas* RNAi[27, 28] in wing discs was achieved using the flip-out system[29]. Larvae from crossing flip122; tub>stop>GAL4, UAS-GFP females to either the wild type strain Or-R (“wild type”) or UAS-gigasRNAi (“gigas”; VDRC stock# 103417) males were heat-shocked at 37°C for 30min at 72-96 hours after egg laying (AEL). Wing discs from crawling third larval stage (≈120h AEL) were dissected, fixed and stained with mouse anti-Armadillo antibody (N27A1, Developmental Studies Hybridoma Bank, University of Iowa; 1/100), which labels the subapical region of epithelial cells, and rabbit anti-GFP (A11122, Molecular Probes; 1/1000). Alexa Fluor 488 and 568 (Molecular Probes) secondary antibodies were used at 1/500. Samples were imaged on a Leica SPE confocal setup, with an oil-immersion 40X objective (N.A.=1,15), as z-stacks of 12 bits, 1024X1024 pixel images, with z-step= 0.3 microns.

### Image Analysis

3D image stacks used in morphometric analyses were first pre-processed using Fiji [30]. We selected image planes that only contained apical cell boundaries as marked by the anti-armadillo antibody staining, removed frames containing non-apical and non-pouch signal, used a maximum intensity projection to project all substacks to 2D, and enhanced their contrast using the Contrast Limited Adaptive Histogram Equalization (CLAHE) [31]. Using Ilastik [32], we then built a random forest classifier that was used for discriminating cell boundaries from background and noise in problematic image regions. The resulting probability maps were binarised and used to build a Voronoi diagram using Fiji. All diagrams (masks) were manually curated using Tissue Analyzer [33] until they exactly mapped to all apical cell boundaries. Pertinent morphometric measurements were derived and exported by using the EpiTools [15] plugin for Icy [34]. The parameters of the Weibull distribution were fitted using the R package fitdistrplus [35]. All plotting was done using several R packages, including ggplot2, gdata, gtools, and data.table [36–42].

### Set-Up of the LBIBCell simulations

The details of how the `LBIBCell` simulations were set up are provided in the Supplementary Text.

## Acknowledgements

We thank Davide Heller and Luis M. Escudero for providing access to their raw data. We are grateful to Anna Stopka and Harold Gómez for advice on `LBIBCell` and image processing. This work has been supported through an SNF Sinergia grant to DI, and grants BFU2012-34324 and BFU2015-66040 (MINECO, Spain) to FC.

## Author Contributions

DI developed the theoretical framework, MK carried out all simulations and created all figures, AI, GV-F, FC contributed the data. DI, FC, MK wrote the manuscript. All authors approved the final manuscript.

## EXTENDED DATA FIGURE

**ED Figure 1:**
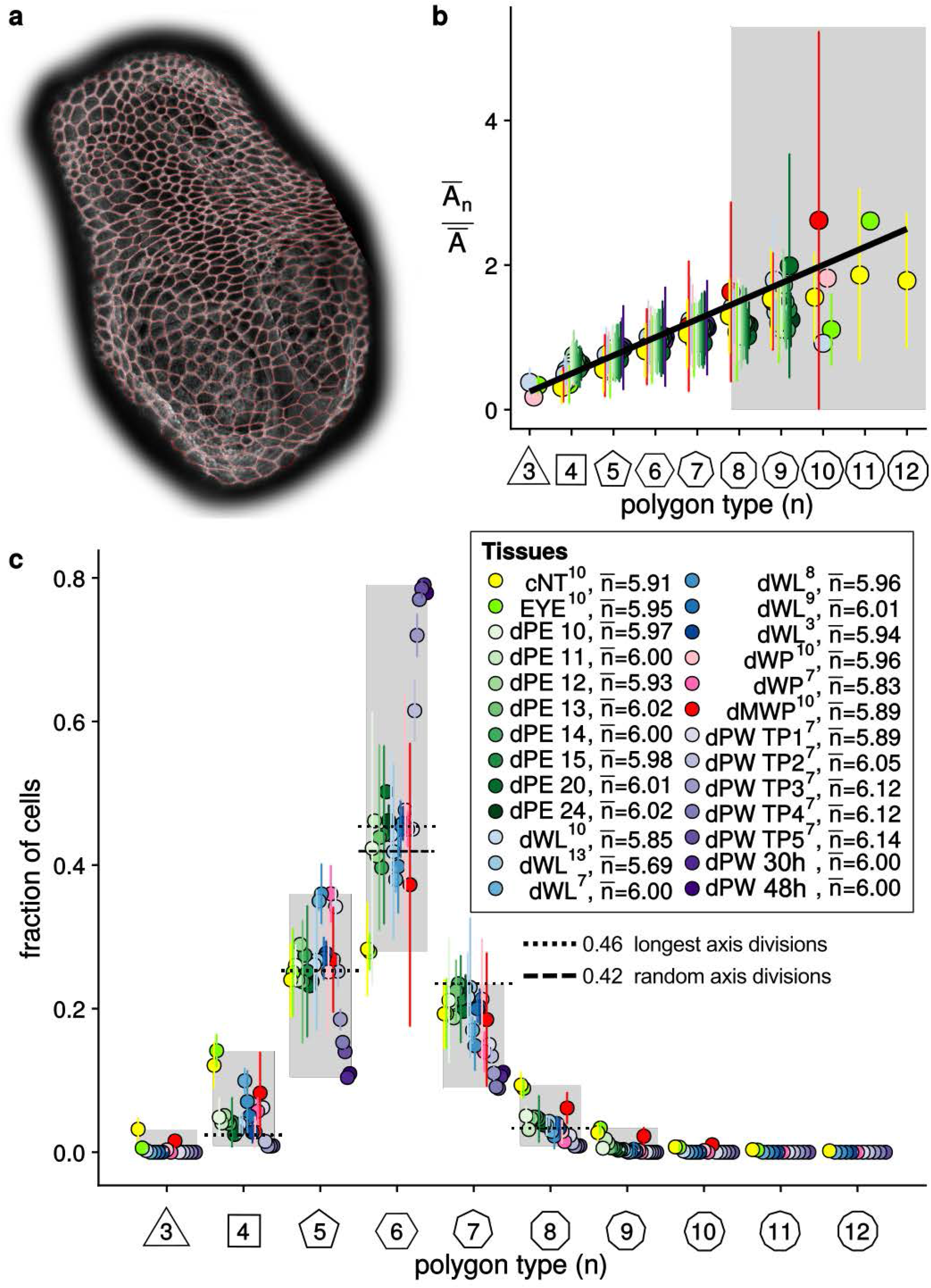
Epithelial Packing in different tissues. **a**, A typical epithelial tissue from the eye disc peripodial epithelium with segmentation mask. **b**, Tissues differ widely in the relative frequency of neighbour numbers. Topological models only explain a single (dashed lines) of these many distributions (shaded range). The average number of cell neighbours is close to the topological requirement [1, 6], 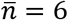, in all tissues; the legend provides the exact values for each tissue and the references to the primary data. The tissue samples are: Chick neural tube epithelium (cNT) [44], *Drosophila* eye disc (EYE) [44], *Drosophila* peripodal membrane from the larval eye disc (dPE), *Drosophila* larval wing disc (dWL) [1, 3, 5, 15, 43–45], *Drosophila* pre-pupal wing disc (dWP) [43, 44], *Drosophila* mutant wing pre-pupa (dMWP: the expression of myosin II was reduced in the wing disc epithelium by expressing an UAS-zipper-RNAi using the C765-Gal4 line) [44], and *Drosophila* pupal wing disc (dPW) [43]. In case of the *Drosophila* peripodal membrane from the larval eye disc (dPE), the numbers indicate the developmental age as the number of ommatidial rows that have formed. In case of the *Drosophila* pupal wing disc (dPW) [44], TPx indicates subsequent, but not further specified pupal time points [43]. Our own measurements of pupal wing discs were at 30 and 48 hours post puparium formation. The datasets dPW and dPE were generated as described in the Methods section; the other datasets were obtained from the specified publications [1, 3, 5, 15, 43–45]. We note that we excluded reported cells with *n* < 3 from the analysis as they must represent segmentation artefacts. **c**, The relative average cell area, 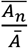, is linearly related to the number of neighbours, n, and follows Lewis’ law[6] (Eq. 2, black line) in all reported tissues[15, 44]. The colour code is as in (b), but data is available only for a subset of those tissues. Few cells with more than 8 neighbours have been measured and estimates of the mean areas are therefore unreliable (shaded part).

**ED Figure 2:**
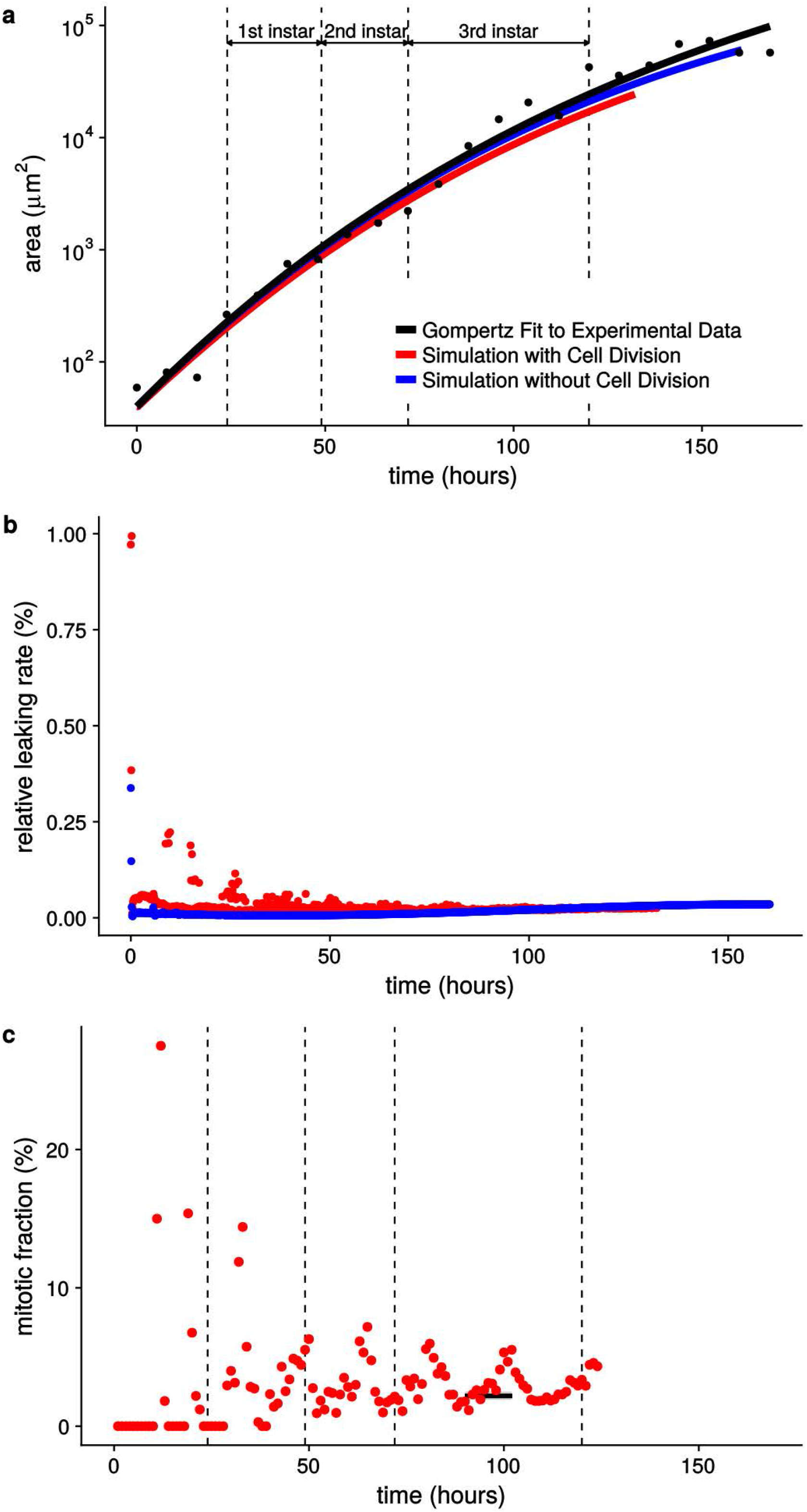
The LBIBCell simulations reproduce the growth dynamics in the Drosophila wing disc. **a**, Simulations with (red) and without (blue) cell division reproduce the measured (black) area growth kinetics[18]. **b**, The small deviation between simulations and measurements is due to numerical leakage (see Supplementary Text for details). **c**, The simulations with cell division (red) reproduce the measured [5] (black) mitotic frequencies in the *Drosophila* larval wing disc. Measurements were reported for mid 3^rd^ larval instar (72-120h AEL). Accordingly, we added the measurements from 84-104h AEL as a black line plus standard deviation in grey.

**ED Figure 3:**
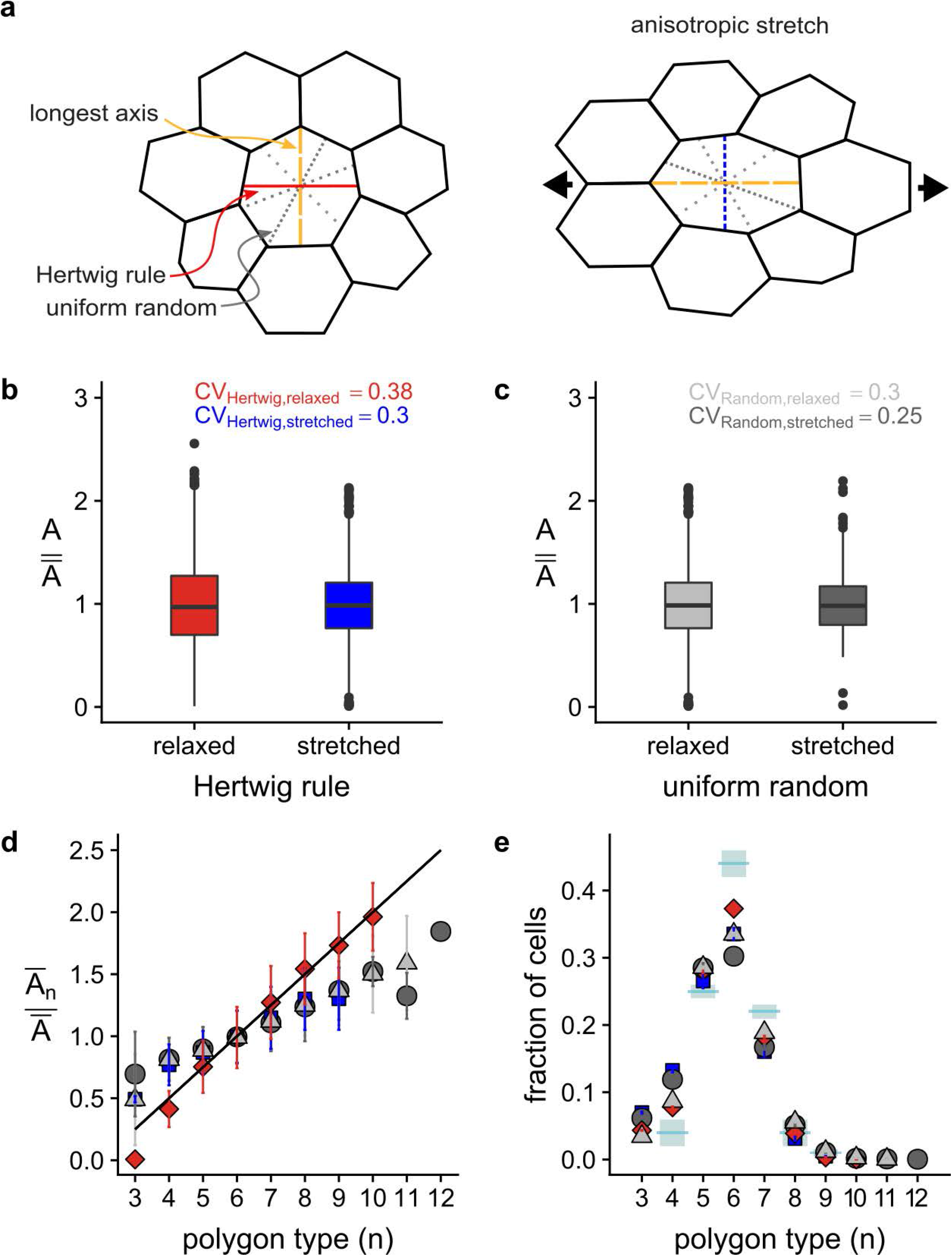
Impact of the Cell Division Axis in LBIBCell. **a**, Cartoon of different cell division axes and of the application of forces to stretch the tissue. According to Hertwig’s rule [7] (red), cells divide perpendicular to their longest axis. For comparison, we simulated a random position of the cell division axis (grey). Moreover, we explored the impact of stretching the tissue in one direction (right hand side). As a result of anisotropic stretching, the cells become elongated and therefore develop a more pronounced longest axis. **b-e**, Impact of the cell division axis (according to Hertwig’s rule (colored) or randomised (grey)) on the cell area distribution (b, c), frequency of neighbour numbers (d), and Lewis’ law (e) in isotropic growing tissue or in anisotropically stretched growing tissue as indicated. Experimental data [15] (light blue) is provided in panel d for comparison.

**ED Figure 4:**
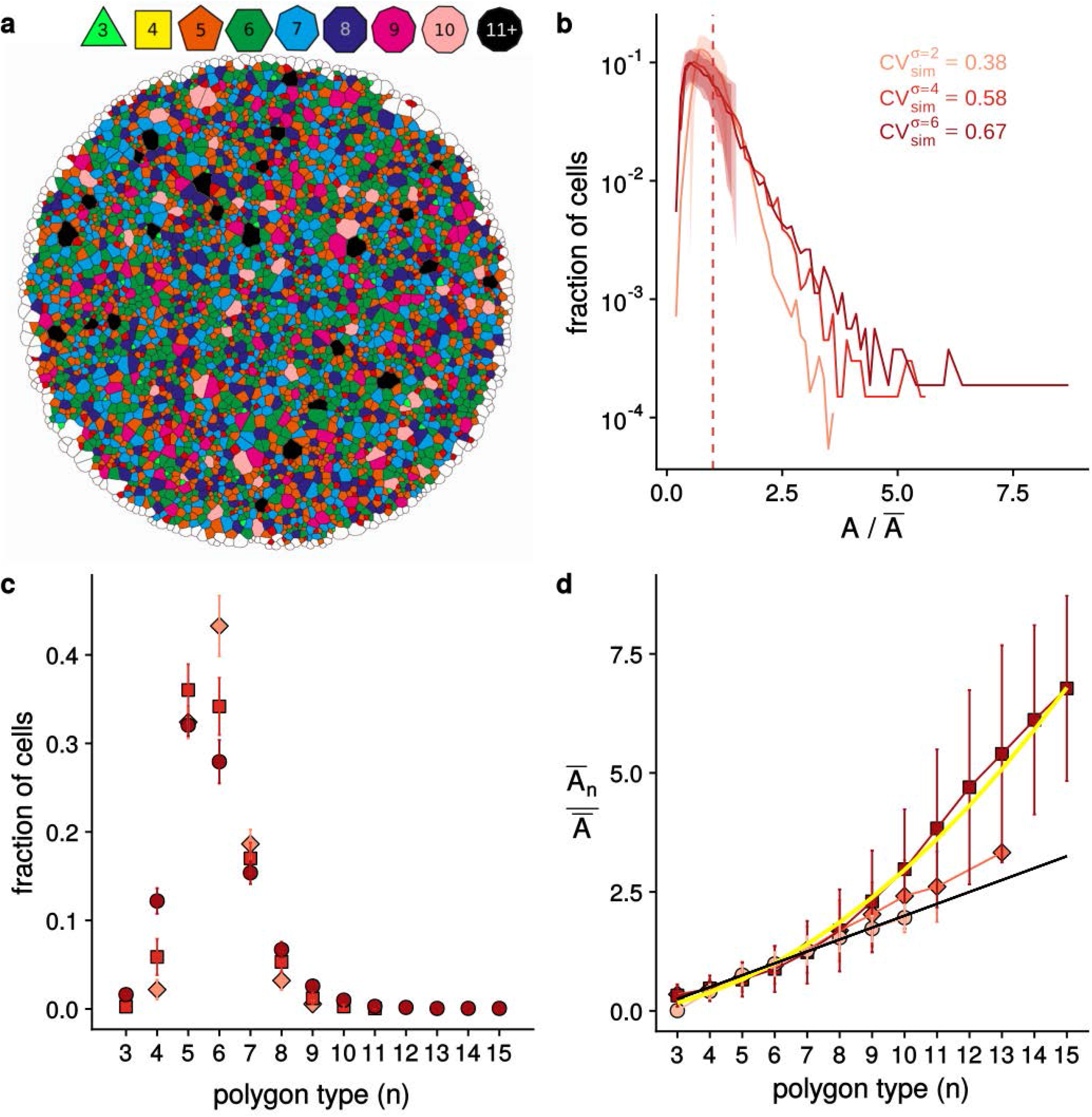
Impact of cell area variability on cell arrangement in LBIBCell. **a**, Representative simulated tissue with high area variability, coloured according to the number of neighbours per cell as indicated. **b**, The cell area distribution and calculated CV value, **c**, the relative frequency of neighbour numbers, and d, the relative average apical area, 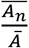 for different neighbour numbers *n*, in simulations with different standard deviations in the cell division threshold (σ=2μm^2^ - light red, σ =4μm^2^ - red, σ =6μm^2^ - dark red); the mean cell division threshold was equal in all simulations (μ=6.7μm^2^). The black line in panel d represents Lewis’ law (Eq. 2); the yellow line the quadratic relation (Eq. 4). Consistent with the theory (Fig. 2c,d), the increase in the cell area variability (panel b) results in a lower fraction of hexagons (panel c).

**ED Figure 5:**
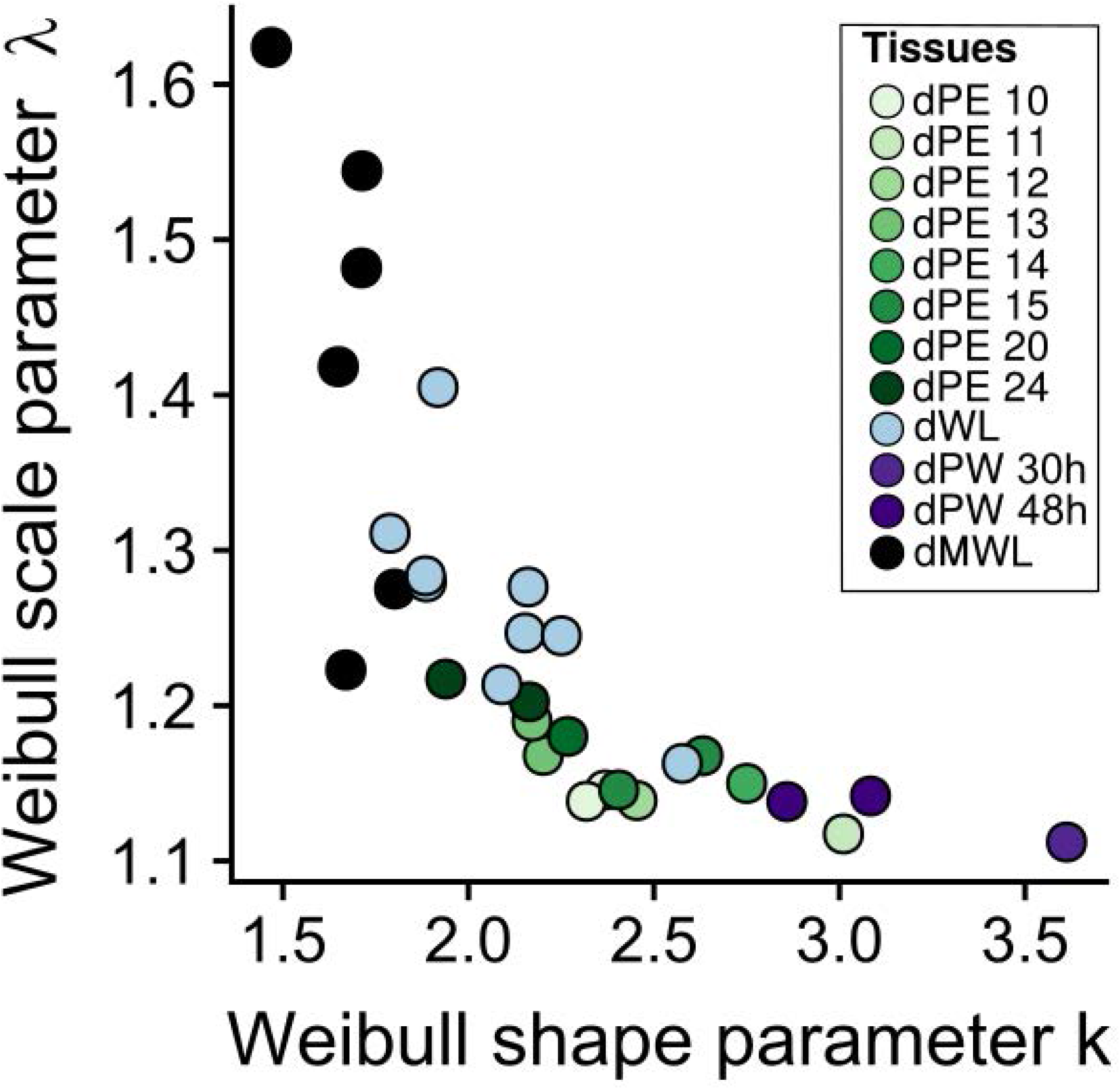
Apical area distribution in epithelial tissues. Weibull (Eq. 6) scale parameter *λ* of the normalized area distribution versus Weibull shape parameter *k* in epithelial tissues. The abbreviations are as in ED Figure 1. dMWL refers to the wing disc with *gigas* RNAi clones from Figure 5.

**ED Figure 6:**
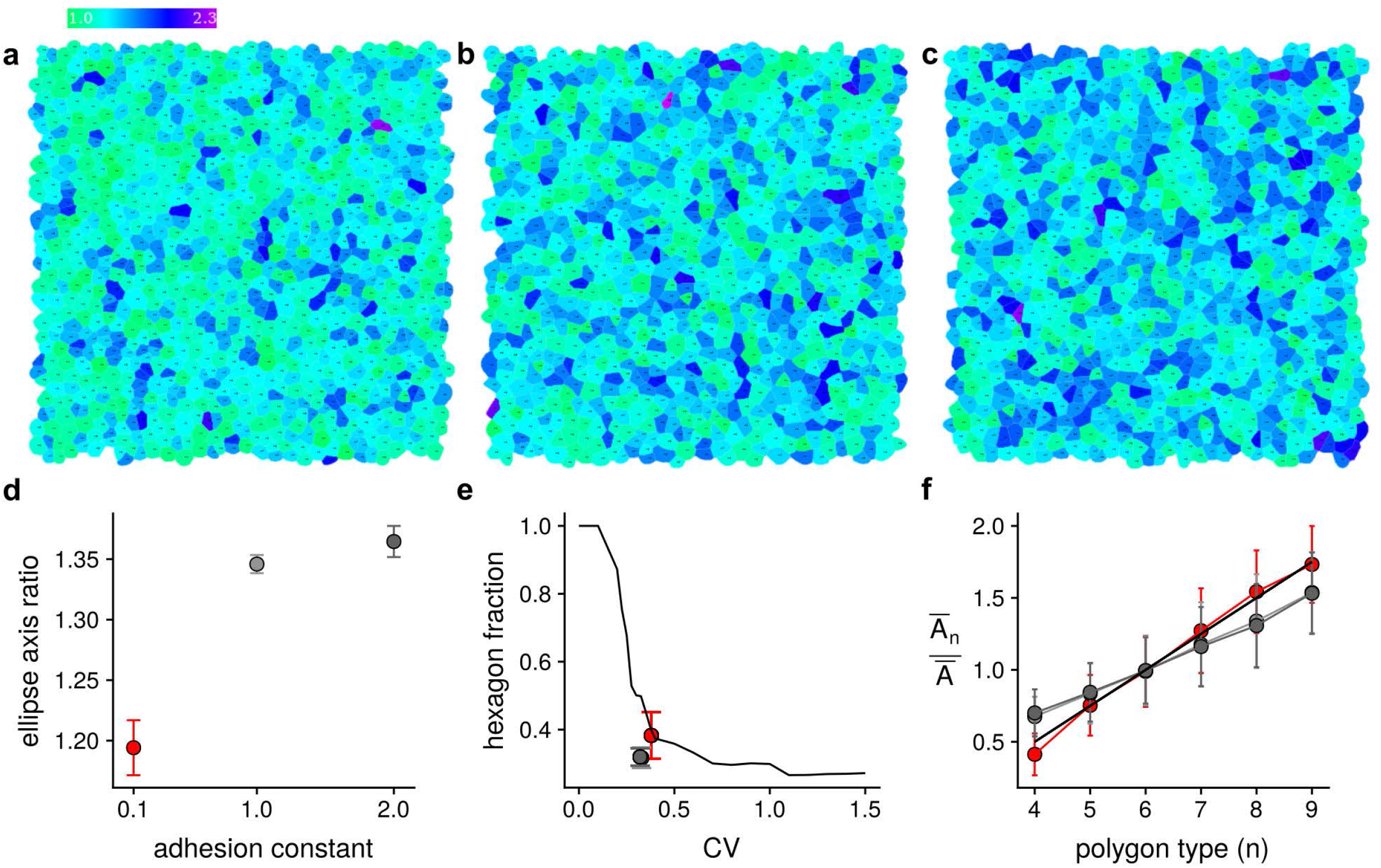
Impact of the relative strength of adhesion strength and membrane tension on Lewis’ law in LBIBCell. **a-c**, Snapshots of the cell lattice in `LBIBCell` for different adhesion constants, i.e. (a) cjK = 0.1, (b) cjK = 1.0, (c) cjK = 2.0. **d**, Impact of the adhesion constant on cell stretching as quantified by determining the average ratio of the long and short axes of ellipses that were fitted to the cells in a simulated tissue for different adhesion constants. **e**, The fraction of hexagons relative to area CV for different adhesion constants. **f**, The relative average cell area, 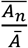, dependent on the number of neighbours, *n*, according to Lewis’ law [6] (Eq. 2, black line) and for `LBIBCell` simulations for different adhesion constants, i.e. cjK = 0.1 (red, standard value as in Figure 4i), cjK = 1.0 (light grey), cjK = 2.0 (dark grey). All other parameter values are as in Figure 4i.

## 1 Supplementary Methods

### 1.1 LBIBCell: Setup of Epithelial Tissue Simulations

We used the software `LBIBCell` [1] to simulate the detailed epithelial tissue dynamics (Movie S1). `LBIBCell` simulates a 2D plane of the epithelial tissue, and thus allows us to simulate the apical cell dynamics, or more precisely the plane where cells adhere laterally (Fig. S1). `LBIBCell` represents the extracellular and cytoplasmic spaces of the tissue as Newtonian fluids and incorporates the fluid-structure interaction with the elastic cell boundaries via an immersed boundary method [2]. The Navier-Stokes equation for the fluid dynamics is solved using the Lattice Boltzmann method, i.e. the fluid and its motion are realized by particle ensembles which stream and collide on a 2D regular lattice [3]. Cell boundaries are geometrically highly resolved and cell boundaries and cellcell junctions are modelled with elastic springs (Fig. S1).

**Figure S1:**
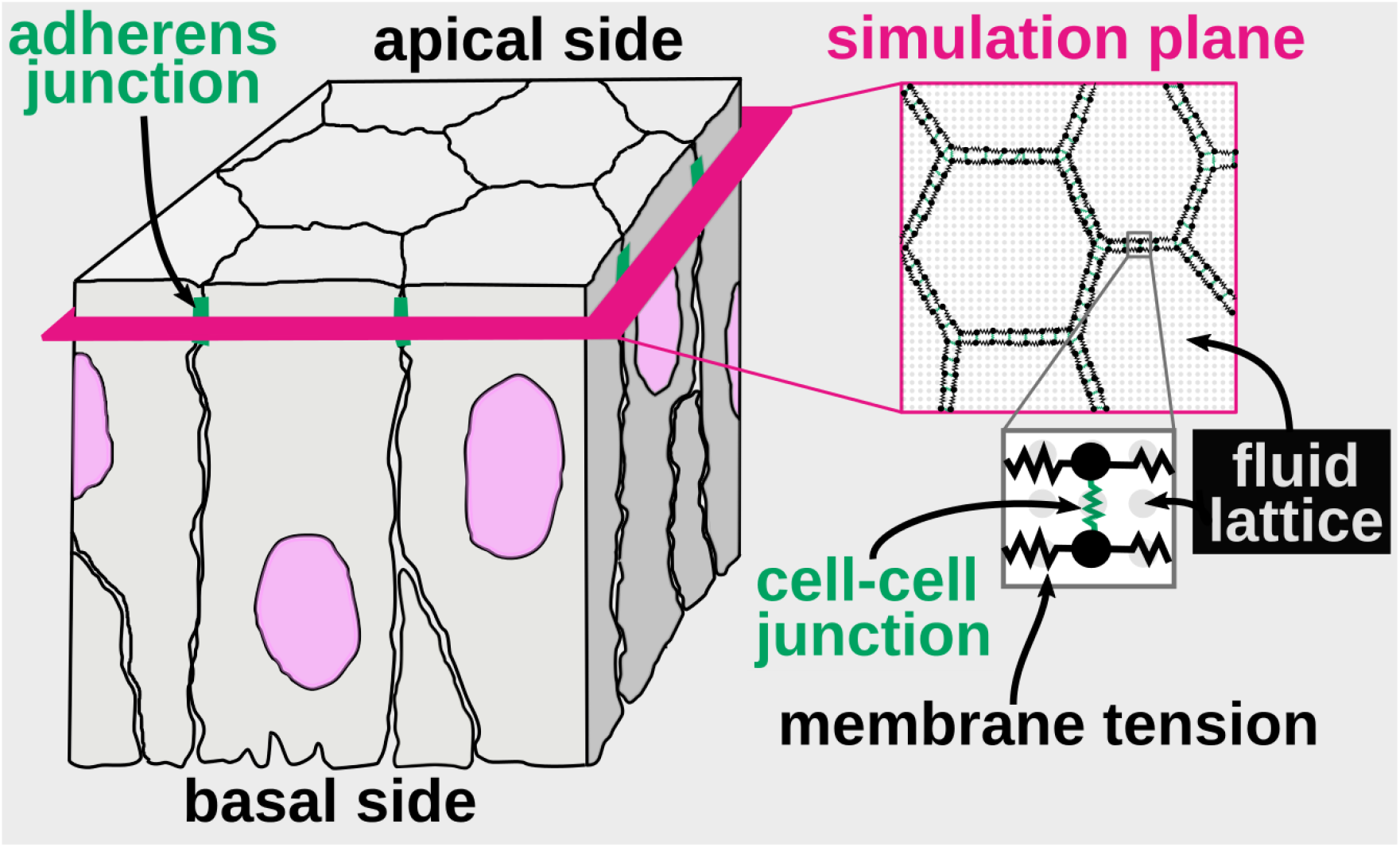
Representation of the apical epithelial surface as a 2D simulation plane in LBIBCell. The cartoon shows a cubical section of a simple columnar pseudostratified epithelium. `LBIBCell` simulates the tissue mechanics and morphodynamics of the 2D plane (indicated in pink) that goes through the adherens junctions (marked in green), which are located below the apical surface. Cell boundaries are geometrically highly resolved and cell boundaries and cell-cell junctions are modelled with elastic springs.

In `LBIBCell`, cellular processes, i.e. cell growth, cell division and cell-cell interactions, are realized by a set of functions, which are implemented in so-called `BioSolvers`. To simulate the epithelial dynamics, we made use of five `BioSolvers` (Table S1). The `Cell Growth BioSolver` expands the cellular domain according to a specified growth rate by adding mass to the intracellular lattice points at each time step *i*. Since the fluid is quasi-incompressible and the density is thus constant, any intracellular mass increase results in an expansion of the cellular domain and thus in cell growth. The `Cell Division BioSolver` divides cells once these exceed a specified cell area, 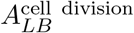, and the `Cell Extrusion BioSolver` removes cells that fall below a specified critical cell area, 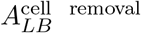. The remaining two solvers, the `Membrane Tension BioSolver` and the `Cell Junction BioSolver` represent the elastic properties of the tissue. The `Membrane Tension BioSolver` describes the cortical tension that is mediated by the acto-myosin ring, while the `Cell Junction BioSolver` models adhesive interactions in the cell-cell-junctions. Both realise linear elastic restoring forces (Hookean springs) between cellular membrane points.

The five `BioSolvers` have a total of 18 parameters (Table S1). Eight of these parameters only affect numerical aspects, and these parameters were set such that an accurate solution was obtained with the least computational effort (Table S1, shaded in grey). The remaining parameters were set based on experimental data. Since the most quantitative data is available for the epithelium in the *Drosophila* larval wing disc pouch, we adjusted the parameter values of the `BioSolvers` to reproduce these measured values. In the following, we describe in detail how the numerical and biological parameter values were set and how their choice was validated.

#### 1.1.1 Numerical Parameters

The numerical parameters do not alter the simulation outcome as long as they lie within a numerically stable range. We therefore chose the numerical parameter values that minimise the computational runtime without affecting the simulation outcome (Fig. S2, Fig. S3).

##### Relaxation time in the Lattice-Boltzmann method

`LBIBCell` uses the standard Lattice Boltzmann scheme (with the single-relaxation-time Bhatnagar-Gross-Krook collision operator) for the fluid [1]. The Boltzmann equation is discretized on a D2Q9 lattice. The only adjustable parameter is the relaxation time *τ*_fluid_ ∈ (0.5,2]. The lower bound *τ*_fluid_ > 0.5 ensures that the kinematic viscosity is larger than zero, i.e 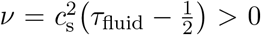. For isothermal flows, the speed of sound is defined as 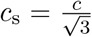 using the lattice speed *c* = Δ*x*/Δ*t*. The relaxation time *τ*_fluid_ thus determines the time discretisation in the Lattice Boltzmann method [1]. For *τ*_fluid_ = 1, the lattice spacing and the time step are Δ*x* = Δ*t* = 1, and are then consistent with the lattice. The upper bound of *τ*_fluid_ results from the stability requirement of the LB-method; in the case of the D2Q9 scheme, *τ*_fluid_ ≤ 2 [4]. We used the highest possible value, *τ*_fluid_ = 2, for maximal computational efficiency (Fig. S2).

**Figure S2:**
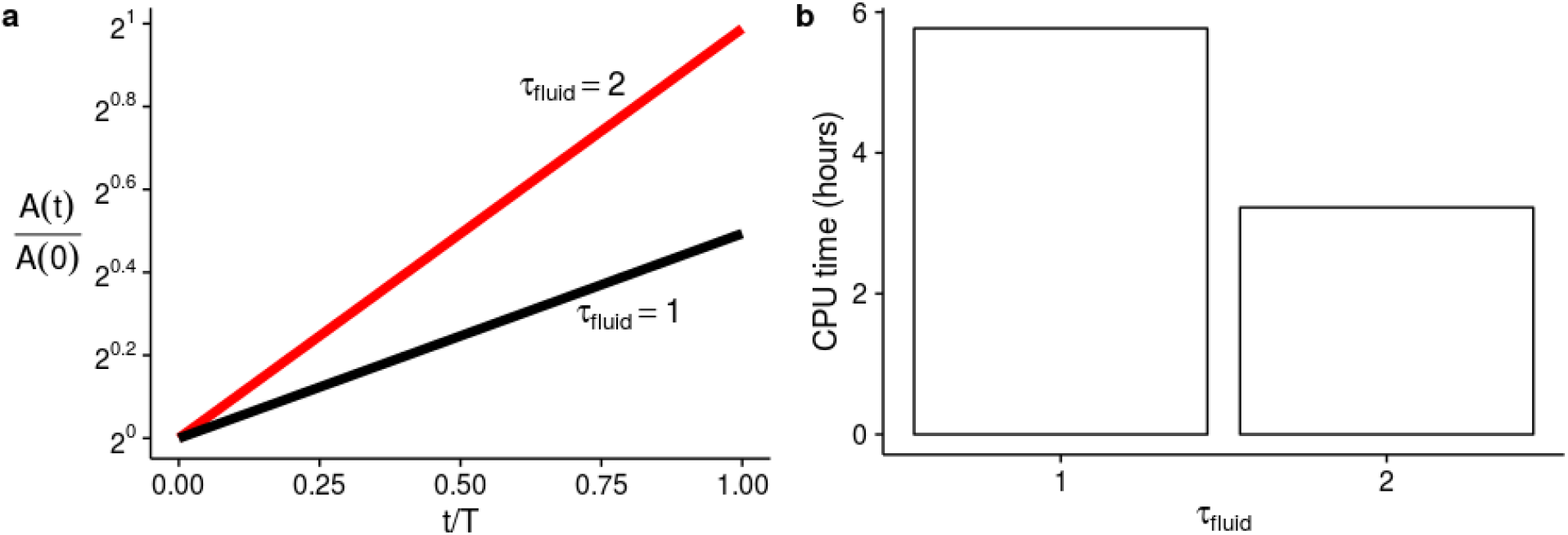
Relaxation time in the Lattice-Boltzmann method. **a**, Growth simulations of a single cell with a constant growth rate *g*(*t*) = 0.5/*T* and *τ*_fluid_ = 1 (black line) or *τ*_fluid_ = 2 (red line). Here, *t* ∈ [0,*T*] is time and *T* is the total time; *A/A*(0) monitors the relative area increase. In LB time units *i* ∈ [0,*I*], we have 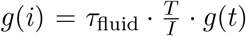, because 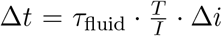. Given the difference in time discretisation, growth per LB time step is double as fast for *τ*_fluid_ = 2 compared to *τ*_fluid_ = 1. **b**, CPU time in hours for a twofold increase in area *A* with *τ*_fluid_ = 1 or *τ*_fluid_ = 2.

##### Lattice resolution

The fluid lattice is regular, and the distance between two lattice points corresponds to 1 LB Length Unit, in short 1*L*_*LB*_. The resolution of the lattice needs to be sufficiently high to avoid numerical artefacts (cf. [1] supplementary and 1.1.3), but otherwise does not affect the simulation outcome. In previous simulations, an absolute minimum of about 100 lattice points per cell was determined [1]. As cells grow and divide in the simulation, we chose the resolution such that even the smallest cells possess at least 100 lattice points. This was realised by setting the area threshold for cell removal (extrusion) to 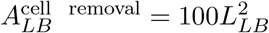. All other area-dependent parameter values were set relative to this number (see below).

**Figure S3:**
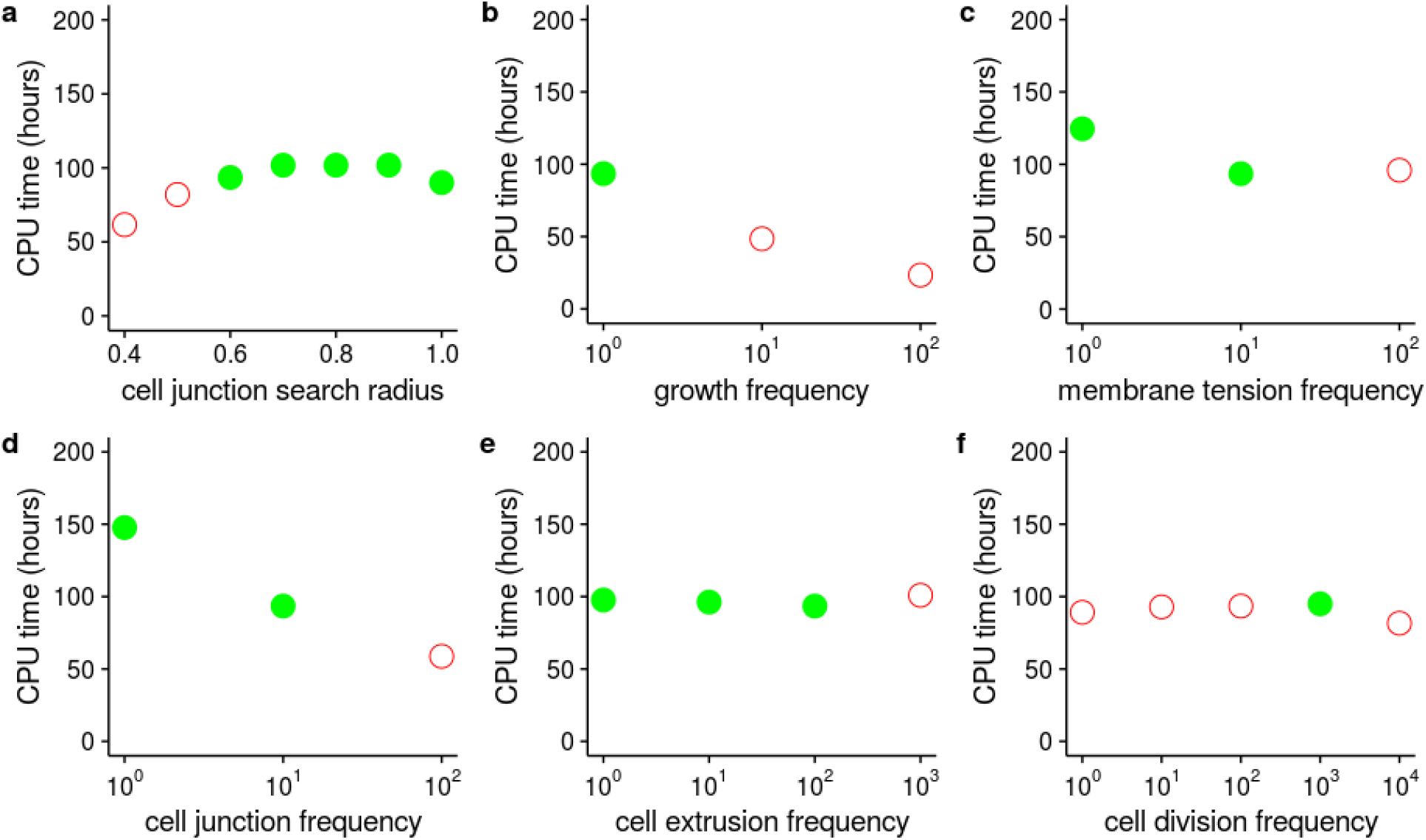
LBIBCell: Numerical parameters for simulations with random cell division threshold. Computational runtime in hours for different choices of the numerical parameters as specified on the the horizontal axes. Numerical parameter values that result in valid tissues without defects, i.e. gaps, overlaps of immersed boundaries or complete disintegration, are depicted by a green filled circle 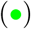. Numerical parameter values that result in invalid tissues, aborted simulations, or insufficient cell numbers in case of the cell division frequency are depicted by a red open circle 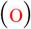. Of the numerical parameter sets that result in valid tissue configurations, the one with the least computational effort was chosen (Table S1).

##### Search radius for new junctions

The search radius *R* for new junctions needs to be in a range that only local, but not distant cell-cell junctions are possible. In particular, the search radius needs to be large enough that new junctions can be established after cell division. During cell division, the mother cell is divided along the cell division axis, and two new membrane segments are introduced as boundaries of the two daughter cells. These have a default distance of 0.5*L_LB_*, which is deliberately larger than the resting length of cell-cell junctions (see below). For too small search radii, gap formation within the tissue occurs. Accordingly, the search radius needs to be at least 0.5*L_LB_*. To avoid connections between distant cells across tissue boundaries, the search radius should be as small as possible. In case of a fixed area threshold for cell division (cf. section on cell division (area) threshold), we can use *R* = 0.5*L_LB_*. In case of a normally distributed area threshold for cell division (cf. section on cell division (area) threshold), larger search radii must be used (Fig. S3a). The exact value depends on the variability in the cell division area threshold; to reproduce the data measured by Heller and co-workers [5], we needed to use at least *R* = 0.6*L_LB_* (Table S1); for consistency, we used *R* = 0.6*L_LB_* also in case of a constant cell division area threshold (Fig. 4d-f in the main manuscript).

##### Frequency of applying the BioSolvers

Five of the eight numerical parameters concern the frequency with which the five `BioSolvers` are applied. The `Cell Growth BioSolver` achieves growth of the domain by adding a specified mass to each lattice point within a cell. Greatest numerical stability is achieved if mass is added in each iteration step (Fig. S3b), and the frequency is thus 1:1 (Table S1). The `Membrane Tension BioSolver` and the `Cell Junction BioSolver` control the forces which act on the cell boundary points (`Geometry Nodes`) and interact with the fluid via the immersed boundary method. To avoid numerical inaccuracies from delayed updates, these solvers have to be called at least every 10th time step (Fig. S3c,d), i.e. 1:10 (Table S1). The `Cell Extrusion` and the `Cell Division BioSolver` remove or divide cells, respectively. Simulation output did not change as long as the `Cell Extrusion BioSolver` was called in every 100th LB step (Fig. S3e). In case of a fixed cell division area threshold (cf. section on cell division (area) threshold), simulation output did not change as long as the `Cell Division BioSolver` was called in every 1000th LB step. In case of a randomly distributed cell division area threshold, it is important to not call the `Cell Division BioSolver` too frequently (cf. section on cell division (area) threshold). We therefore called the `Cell Division BioSolver` in every 1000th LB step in all simulations (Fig. S3f).

#### 1.1.2 Biological Parameters

The biological parameter values had to be adapted such that the simulated tissue matches the properties of the *Drosophila* larval wing disc pouch epithelium.

##### Conversion between LB units and SI units

LB units can be converted to metric units based on the simulated developmental time periods and the measured cell areas. Thus, in the simulations, time is discretised as *i/I* ∈ [0, 1], where *i* refers to the iteration step and / to the total number of iterations. Each iteration step *i* thus corresponds to Δ*t* = *τ*_fluid_ · *T/I* hours, where *T* is the total simulated developmental time. For restrictions on the choice of *I*, see the section on *Numerical Leakage at the Fluid-Structure Interface*.

The smallest measured apical cell area is 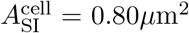, the largest measured apical area of non-dividing cells is 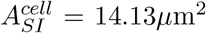, and the average apical cell area is *Ā*_SI_ = 5.34*μ*m^2^ [5]. As we set the threshold for cell removal to 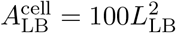, the conversion factor between metric units and LB units follows as

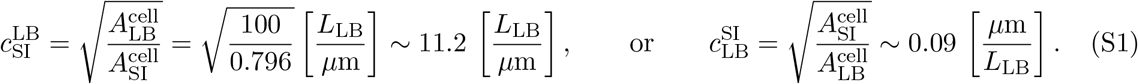

##### Elastic Tissue Properties

The `Membrane Tension BioSolver` and the `Cell Junction BioSolver` describe the elastic properties of the tissue. The membrane tension (Fig. S4a) represents the cortical tension mediated by the acto-myosin ring and was realised by adjusting the magnitude *φ^m^* of a constant contracting force, 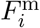, between pairs of cell boundary points with position vectors **x** (Fig. S4a), i.e.

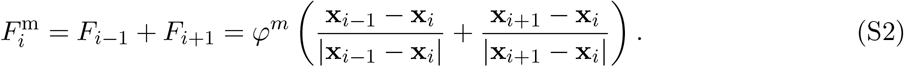

In the case of the cell-cell junctions (Fig. S4b), a Hookean force 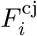 with a spring constant *k_j_* and resting length *l*_0_ acts between intercellular cell boundary points *i* and *j*,

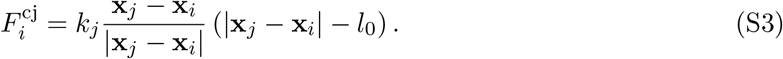

The resting length *l*_0_ was set to the length of cadherin junctions, which has been reported as 26-39nm, depending on the type of the bond [6]. We used 0.3*L_LB_*, which corresponds to 27nm. This value is deliberately smaller than the default distance between sister cells after cell division of 0.5*L_LB_* = 45nm (see above), which is on the high end of distances between boundaries of neighbouring epithelial cells.

The choice of spring constants determines the neighbour frequencies in the simulated tissue, i.e. larger spring constants lead to an increase in the fraction of hexagons (Fig. 4b in the main manuscript). For very large spring constants, cells become rounded, while for very small values, cells adapt an angular shape and, in combination with a low cell junction spring constant, the cell boundaries become curled. We adjusted the spring constants *φ*^m^ and *k_j_* to reproduce the measured neighbour frequencies [5] (Table S1).

**Figure S4:**
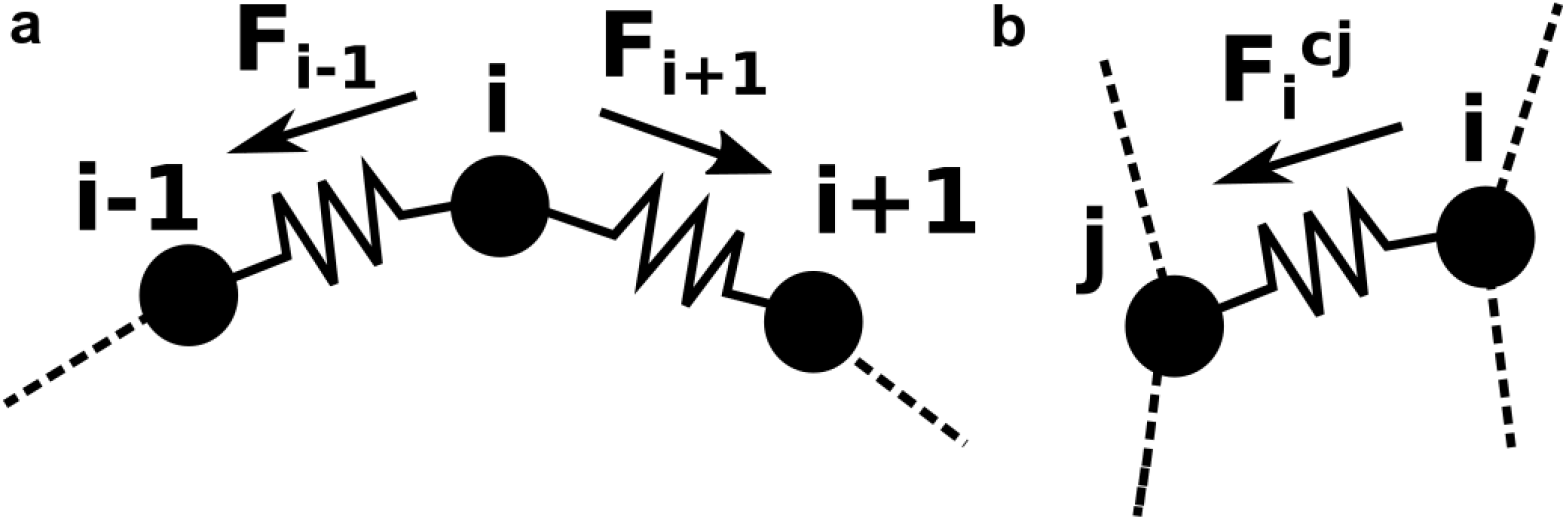
Hookean springs to model Elastic Tissue Properties. **a** Membrane tension: the superposition of Hookean spring forces *F*_*i*−1_, *F*_*i*+1_ on a cell boundary point *i* results in a contracting force 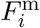. **b** Cell-cell junctions: the force 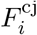 depends on the spring constant *k_j_* and the deviation of the distance between the two cell boundary points x_*i*_ and x_*j*_ from the resting length *l*_0_.

**Figure S5:**
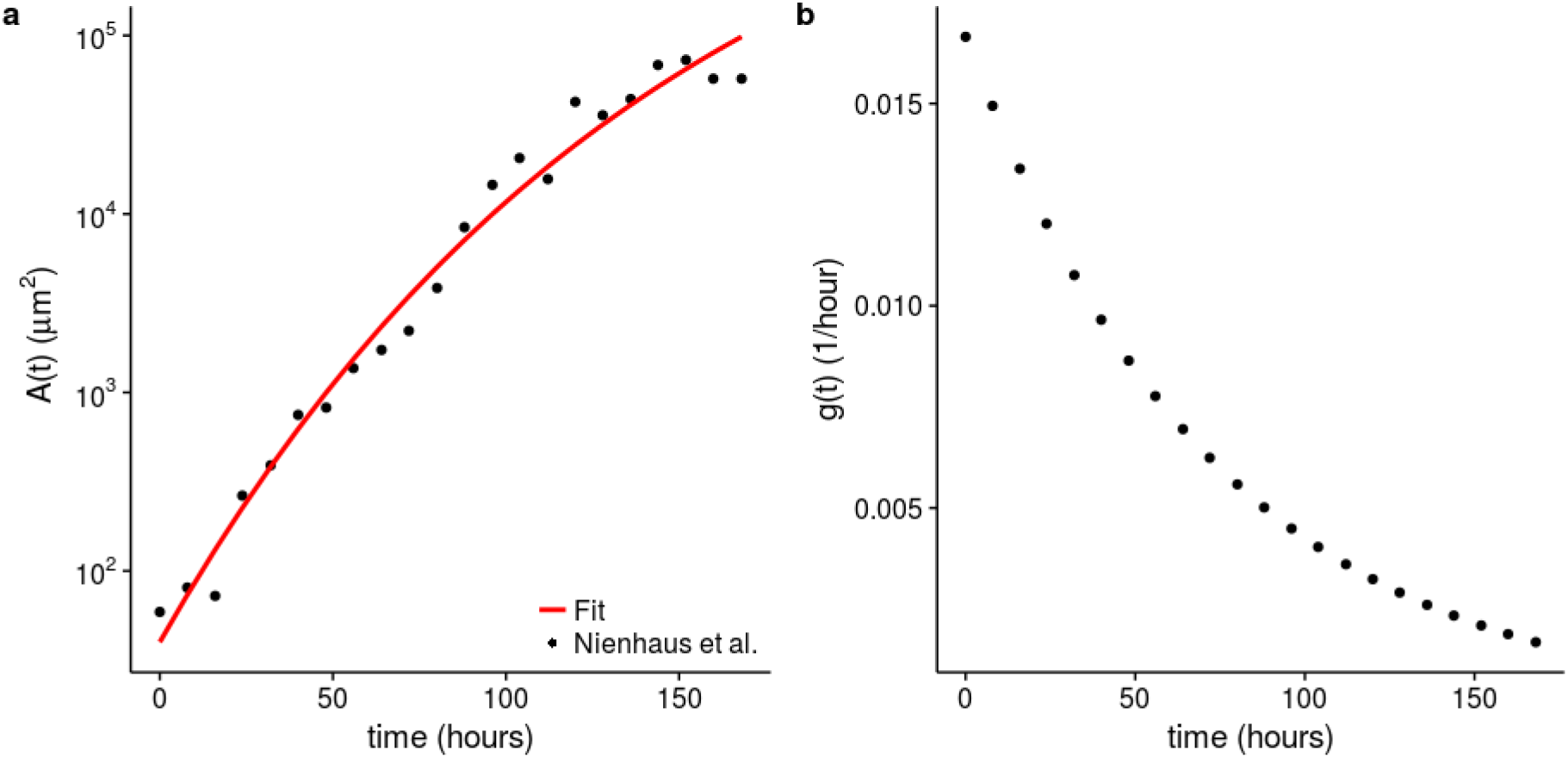
Inference of the apical area growth rate from published area growth data from the *Drosophila* larval wing disc. **a**, Fit (red line) of the apical area growth data (black dots) from the *Drosophila* larval wing disc [7] with Gompertz function (Eq. S4). **b**, Area growth rate *g*(*t*) according to equation S5 with the parameters of the fit in panel a.

##### Growth Rate

We sought to reproduce the measured area growth rate in the *Drosophila* larval wing disc pouch [7]. As shown before [8,9], the area growth data is fitted well (*R*^2^ = 0.98) by Gompertz growth kinetics of the form

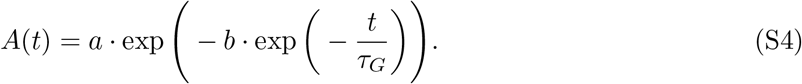

with *a* = 3.327 · 10^6^ *μm*^2^, *b* = 11.331, *τ_G_* = 143.777 hours (Fig. S5a). The corresponding growth rate that was used in the simulations is thus an exponentially declining function (Fig. S5b),

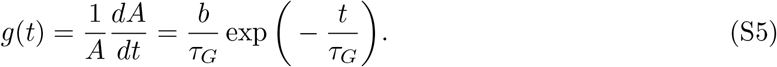

In LB units, we then have

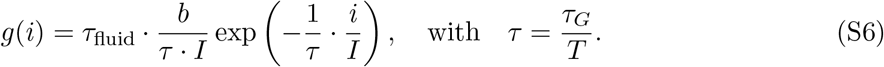

##### Cell division threshold

The `Cell Division BioSolver` uses the cell area as a threshold for the initiation of cell division. We implemented two different cell division thresholds. In the first case (shown in Fig. 4d-f in the main manuscript), all cells were divided when reaching a fixed critical cell size. The critical cell size was set to 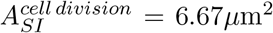 such that the measured average cell area of 5.34*μ*m^2^ [5] was reproduced. In LB units, 6.67*μ*m^2^ correspond to 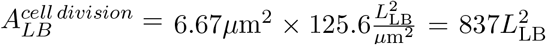. In the other cases, the cell division threshold followed a normal distribution with mean 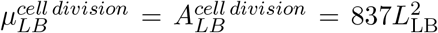 and standard deviation as specified in the respective panels, i.e. see Fig. 4g,j and ED Fig. 4 in the main manuscript. In particular, to reproduce the data by Heller and co-workers [5], we used as standard deviation 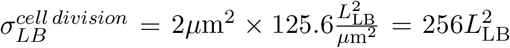. To pick cells randomly according to these distributions, we used a pseudorandom number generator based on the Mersenne Twister algorithm [10]). In addition to the cells that exceeded the randomly picked cell division threshold also those cells were divided whose area exceeded the largest measured apical cell area in the wing disc (14.13*μ*m^2^) [5], i.e. 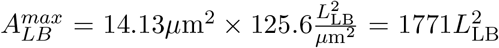. As before, the `Cell Division BioSolver` was applied in every 1000th simulation, but only to 33% of the cells to enable some cells to grow to a larger size before being divided.

##### Cell division plane

According to Hertwig’s rule, cells divide perpendicular to their longest axis [11,12,13]. Accordingly, in the `Cell Division BioSolver`, cells were divided perpendicular to the longest axis of the cell. We also explored the effect of a random orientation of the cell division axis, but the effects were found to be minor, even when cells are stretched (ED Fig. 3).

##### Initial Conditions

*Drosophila* larval wing discs are derived from 22-34 precursor cells (> 95% confidence interval) that give rise to about 30000 cells at the end of the larval stage [14]. Thus, when simulating the entire developmental phase of the larval wing disc pouch (ED Fig. 2a), we start our simulations with a tissue patch of *n* = 2^5^ = 32 cells. According to the fit to the larval wing disc area growth data [7], the initial larval wing disc area at 0 hours after egg lay (h AEL) is 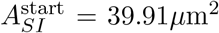 (Fig. S5a). According to Eq. S1, 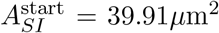 corresponds to 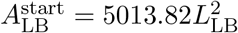 in LB units.

Most experimental studies of the epithelial organisation of *Drosophila* larval wing discs focus on a subset of the mid-third larval instar pouch tissue, and the study we use reported 10^2^−10^3^ cells per tissue sample [5]. When comparing our simulations to this data (Fig. 4 in the main manuscript), we simulated a patch growing to a similar final size in `LBIBCell`. To this end, we started with a tissue patch of *n* = 32 cells. The total initial area was set to 225.40*μ*m^2^, corresponding to the fit to the data [7] at 24 h AEL. According to Eq. S1, 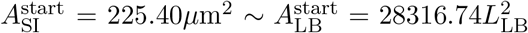. We then simulated the growth process from *t* = 24 h AEL until *t* = 96 h AEL, representing the time period from early first to mid-third larval instar. All results reported in Fig. 4 and ED Fig. 4 in the main manuscript were quantified at the end of this simulation period.

#### 1.1.3 Numerical Leakage at the Fluid-Structure Interface

Due to the discretized nature of both the fluid and the immersed boundaries, fluid leaking at the fluid-structure interface can occur [2]. This numerical leaking, or lack of volume conservation, is caused by a combination of excessive forces acting on the immersed boundary points and the interpolation method of the fluid velocity close to the moving immersed boundary, and is a well-known limitation of the discrete immersed boundary method [1].

The relative leakage rate was calculated as the relative error *δ_A_* between the fitted measured values and the simulation values of subsequent timesteps with a stepping size of 100 iterations: 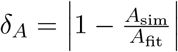.

Contributing factors are both the fluid source strength, i.e. the growth rate *g*(*i*) in each iteration step *i*, and the tension at the immersed boundary. The absolute value of the growth rate *g*(*i*) depends on the total number of iterations *I*, and we used *I* = 150000 to minimize leakage for our specific biological parameters (Fig. S6). For higher values of *I* simulations aborted.

**Figure S6:**
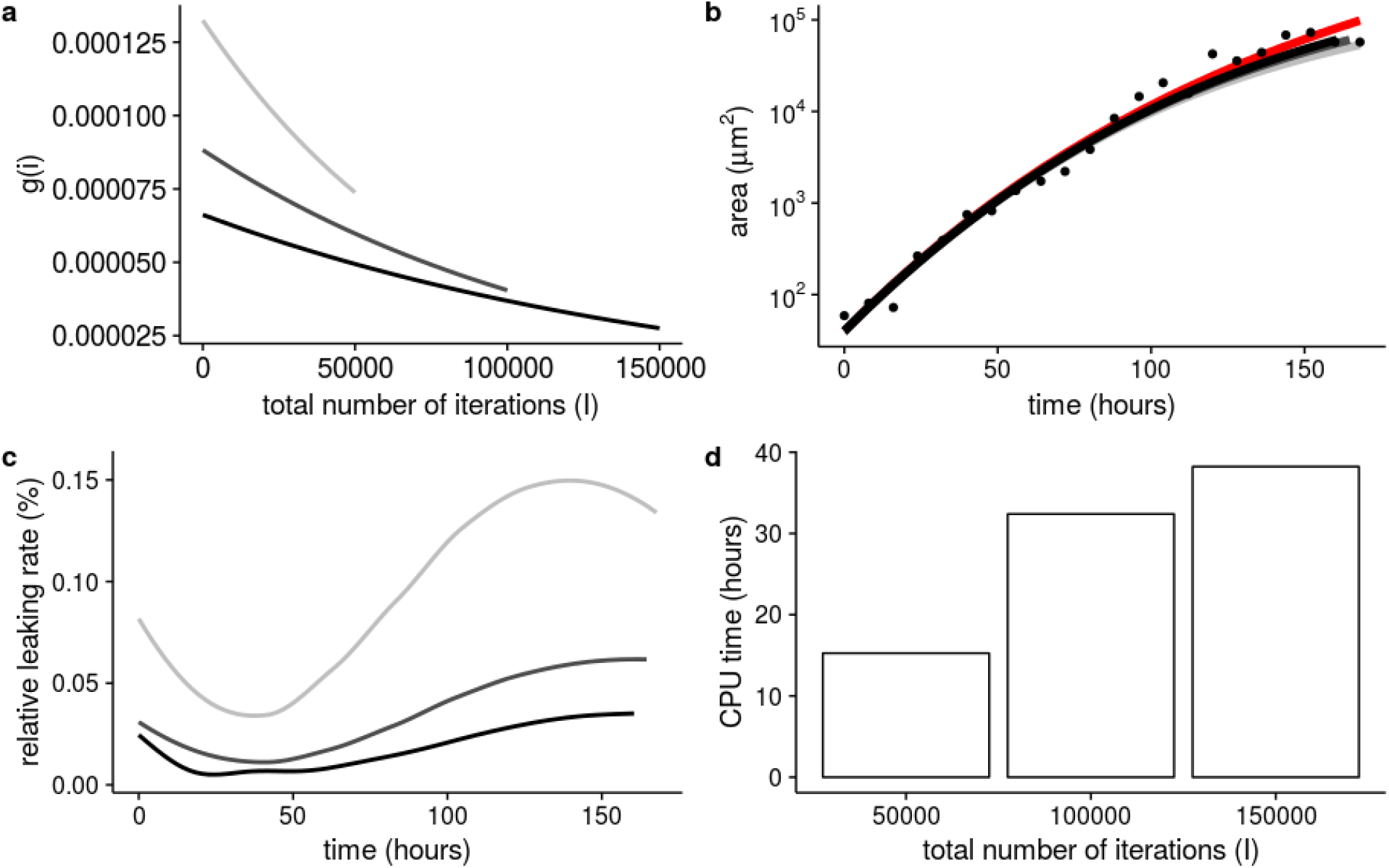
Impact of the total number of iterations *I* on numerical leaking. **a**, Growth rate *g*(*i*) according to Eq. S6, **b**, area growth, and **c**, leakage rates for *I* = 50000 (light grey), *I* = 100000 (dark grey), and *I* = 150000 (black). The red line in panel b represents the fit to the experimental data [7] (dots) as in Fig. S5a. **d**, CPU times for the different *I*.

## 2 Supplementary Tables

**Table 1:** Default parameter values. Parameters in grey-highlighted rows only affect the numerical accuracy of the simulations. The other parameter values are specific to the biological problem of interest, and were determined based on published quantitative data for the *Drosophila* larval wing disc pouch. Here, the parameter values resulting in the best match to the data are provided; the corresponding simulation results are shown in Fig. 4g-i in the main manuscript. Deviations from these default parameter values are indicated in the respective figure legends.

|  |  |  |  |  |
| --- | --- | --- | --- | --- |
| **LBSolver** | Parameter Description<br>$\tau_{\text{fluid}}$ - relaxation time | Value [LB units]<br>2 | Value [SI units]<br>N/A | Determination / Literature<br>Fig. S2, [4] |
| **BioSolver** | Parameter Description | Value [LB units] | Value [SI units] | Determination / Literature |
| **Cell Growth** | frequency of applying BioSolver | 1:1 | N/A | Fig. S3b |
| | $I$ - total number of iterations | 64300 / 150000 | 24-96h AEL / 0-168h AEL | Fig. S6 |
| | $b$ - Gompertz constant | 11.331 | 11.331 | Fig. S5a / [7] |
| | $\tau$ - Gompertz constant | 0.8555 | 143.7773h/168h | Fig. S5a / [7] |
| **Membrane Tension** | frequency of applying BioSolver | 1:10 | N/A | Fig. S3c |
|  | Hookean spring constant | 0.01 | N/A | Fig. 4b / [5] |
| **Cell Junction** | frequency of applying BioSolver | 1:10 | N/A | Fig. S3d |
|  | search radius for junction building | 0.6 | N/A | Fig. S3a |
|  | length of intercellular cell junction | 0.3 | 27nm | 16-39nm [15] |
|  | Hookean spring constant | 0.1 | N/A | Fig. 4b / [5] |
| **Cell Division** | frequency of applying BioSolver | 1:1000 | N/A | Fig. S3f |
| | maximal cell area | 1771 | $14.1\mu\text{m}^2$ | [5] |
| | mean cell division threshold | 837 | $7\mu\text{m}^2$ | Fig. 4g / [5] |
| | std cell division threshold | 256 | $2\mu\text{m}^2$ | Fig. 4g / [5] |
| | orientation of cell division plane | $\perp$ to longest axis | $\perp$ to longest axis | Hertwig's rule, [12] |
| **Cell Extrusion** | frequency of applying BioSolver | 1:100 | N/A | Fig. S3e |
| | minimal cell area | 100 | $0.8\mu\text{m}^2$ | [1, 5] |

## 3 Supplementary Video Legends

**Movie S1: `LBIBCell` - Cell shape dynamics in a growing tissue.** The movie shows an example of a simulated tissue. The different polygon types are marked by colour; polygons with more than nine neighbours and polygons that are not entirely surrounded by other cells, i.e. boundary cells, are marked in black.

